# Ulacamten: A Novel, RLC-Targeting Cardiac Myosin Inhibitor for Potential Treatment of Cardiac Hypercontractility, Including HFpEF

**DOI:** 10.64898/2026.01.23.701387

**Authors:** Saswata Sankar Sarkar, Meredith A. Redd, James J. Hartman, Darren T. Hwee, Anand Bat-Erdene, Leo Kim, Chihyuan Chuang, Cassady Rupert, Najah Abi-Gerges, Janette Rodriguez, Desirae Martin, Andre deRosier, Samantha Edell, Yangsong Wu, Lisette Yco, Anne N. Murphy, Bradley P. Morgan, Fady I. Malik

## Abstract

**Background:** Cardiac myosin inhibitors (CMIs) demonstrate advantages over other guideline-directed therapy for patients with obstructive hypertrophic cardiomyopathy (oHCM). By reducing hypercontractility, CMIs abrogate excessive systolic function and improve diastolic function; diminish hypertrophy of the left ventricle (LV); and improve exercise capacity, functional class, and symptoms. Whether CMIs are therapeutic in heart failure with preserved ejection fraction (HFpEF) is of interest because a significant subset of these patients demonstrate supranormal ejection fractions and abnormal LV structure, characteristics in common with HCM, where CMIs have proved effective.

**Objectives:** Our goal was to characterize the mechanism of myosin inhibition for ulacamten and determine its efficacy in a rodent model of HFpEF.

**Methods:** Ulacamten was characterized using biophysical and biochemical approaches, cardiomyocytes from humans and the ZSF1 obese rat model of HFpEF, hypercontractile human-engineered heart tissues, and echocardiography in the ZSF1 rat model.

**Results:** Unlike the other CMIs, aficamten and mavacamten, ulacamten binds outside the S1 domain of myosin and requires the regulatory light chain domain to bind and inhibit the activity of 2-headed myosin. Ulacamten only partially inhibits the myosin ATPase activity in both myofibrillar and protein systems, but inhibition of contractility was nearly complete in cardiomyocytes. Improvement in relaxation was demonstrated in hypercontractile-engineered heart tissues, and chronic treatment of ZSF1 obese rats showed benefits in both cardiac structure and function.

**Conclusions:** Ulacamten inhibits myosin in a manner distinct from aficamten and mavacamten, potentially broadening the mechanistic properties of CMIs available for treatment of hypercontractile cardiac dysfunction.

**CONDENSED ABSTRACT:** Cardiac myosin inhibitors (CMIs) abrogate excessive systolic function and improve diastolic function, diminish cardiac hypertrophy, and improve exercise capacity in humans with obstructive hypertrophic cardiomyopathy (oHCM). Supranormal ejection fraction underlies heart failure with preserved ejection fraction (HFpEF) in some patients. We describe a new CMI, ulacamten, with binding and inhibitory properties distinct from two other FDA-approved CMIs, aficamten and mavacamten. Specifically, ulacamten requires 2-headed myosin to inhibit activity, whereas aficamten and mavacamten inhibit single-headed myosin. Ulacamten inhibits contractility in primary myocytes isolated from control human and hypercontractile ZSF1 obese rat hearts, as well as engineered heart tissues created with induced pluripotent stem cell cardiomyocytes bearing an HCM mutation. Chronic treatment of ZSF1 obese rats as a preclinical model of HFpEF improves diastolic function and reduces hypertrophy and fibrosis, broadening the potential mechanistic landscape of CMIs.

**Visual abstract:** 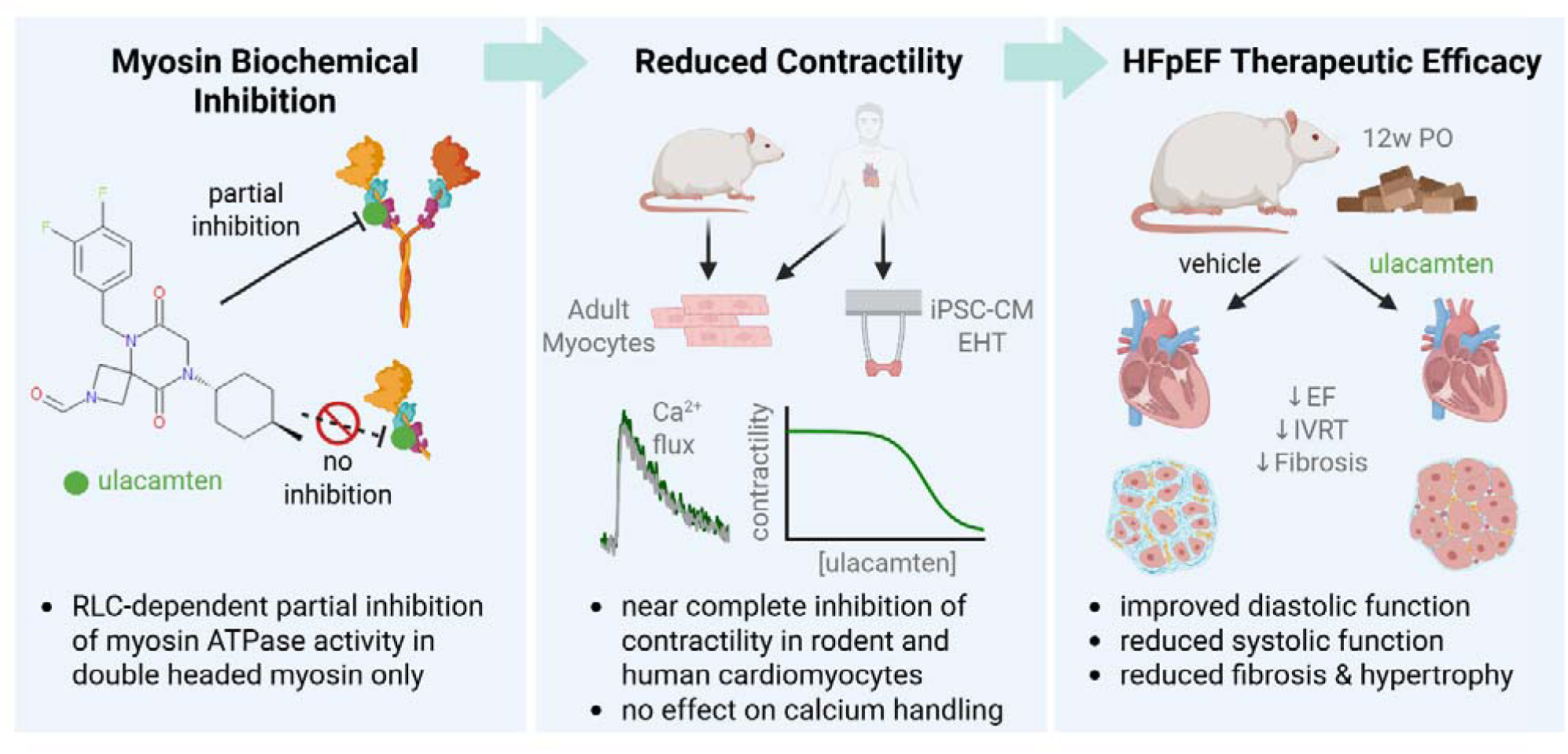

## INTRODUCTION

Heart failure is a leading cause of death and an economic burden to the healthcare system. Patients with heart failure are clinically divided into 3 subgroups based on the contractile phenotype of cardiac function: heart failure with reduced ejection fraction (HFrEF), heart failure with mildly reduced ejection fraction (HFmrEF), and heart failure with preserved ejection fraction (HFpEF).^1^ Patients with HFrEF or HFmrEF suffer from hypocontractility of the left ventricle (LV) with ejection fraction (EF) <50%, whereas patients with HFpEF retain normal contractile LV function (≥50% EF). To date, only sodium-glucose co-transporter 2 (SGLT2) inhibitors have been documented to show clear clinical benefits among heart failure patients with sub-and supranormal EF (EF ≥65%).^2–4^ Although approved for treating patients with HFrEF, beta-blockers are reported to have adverse effects on HFpEF patient outcomes.^5^

Modulation of cardiac contractility with small molecules to treat patients living with heart diseases has attracted much attention in the past 2 decades. The use of omecamtiv mecarbil to increase cardiac contractility in patients with HFrEF proved to be effective in reducing heart failure hospitalizations in a large, randomized phase 3 heart failure clinical trial.^6^ On the other hand, the inhibition of hypercontractile LV function with cardiac myosin inhibitors (CMIs) has proven effective in treating patients with obstructive hypertrophic cardiomyopathy (oHCM).^7^ These patients, who predominantly develop supranormal LVEF, have obstruction of blood flow out of the LV chamber that limits their regular physical activity. Normalizing cardiac contractility with CMIs such as mavacamten and aficamten has proven to be effective in improving peak oxygen consumption, leading to significant improvement in the quality of life and health status of patients with oHCM.^8^ A trend toward the reversal of a thickened LV wall of patients with oHCM is another beneficial outcome of CMI treatment.^9–11^ Furthermore, when tested head to head, each as monotherapy, aficamten was superior to the beta-blocker metoprolol in improving peak oxygen consumption, leading to significant improvement in the quality of life and health status of patients with oHCM,^12^ as well as demonstrating improvement in measures of LV diastolic function, mitral valve systolic anterior motion, and mitral regurgitation in patients with oHCM.^13^

It is increasingly appreciated that the success of CMIs in clinical practice depends on the drug’s safety profile, largely driven by the pharmacokinetic–pharmacodynamic properties, including a shallow dose-response curve, ready reversibility, appropriate half-life, and avoidance of drug–drug interactions.^14,15^ The effectiveness of current CMIs provides incentive to identify additional mechanisms of action that are differentiated from existing ones and perhaps lead to a superior therapeutic profile in humans in HCM and potentially HFpEF. When mavacamten was tested among patients with non-obstructive HCM (nHCM), there was an improvement in clinical biomarkers without significant improvement in health status^16,17^

Thus, we sought to identify inhibitors of sarcomere function with a mechanism of action distinct from aficamten or mavacamten. Here we describe the first such cardiac myosin inhibitor, ulacamten, using a combination of in vitro and in vivo studies.

## METHODS

### Ethical Approvals

Experiments using animals were performed at Cytokinetics using protocols and procedures approved by the Cytokinetics Institutional Animal Care and Use Committee.

### Purification of Myofibrils

Purification of myofibrils was done as reported previously.^18^ Three different isoforms of myofibrils—bovine cardiac, bovine masseter (slow skeletal), and rabbit psoas (fast skeletal) tissue—were prepared from flash-frozen muscle as described previously.^19^

### Purification of Proteins

Bovine cardiac subfragment-1 (S1) and heavy meromyosin (HMM) were prepared as performed previously.^18,20^ Chicken gizzard smooth muscle myosin was purified following the procedure as described previously.^21^ Ultracentrifugation (Beckman Type 45Ti rotor; 142k *g*max, 2.5 h, 4 °C) was performed to clarify protein followed by the addition of 10% sucrose (w/v). Protein solution was drop frozen in liquid nitrogen before storing at −80 °C. Concentration of S1 was determined by measuring absorbance in 6M guanidine hydrochloride using an extinction coefficient of 0.81 cm^2^ mg^-1^. Preparation of bovine cardiac actin from the LV acetone powder of bovine heart was described previously. Actin concentrations were determined by measuring absorbance at 290 nm using 6M guanidine hydrochloride and an extinction coefficient of 6.3 for a 1% solution. The amount of adenosine triphosphate (ATP) in buffer was considered for the measurement of actin concentration.

### ATPase Assays

Steady-state activity measurements of myofibrils and actin-activated ATPase of myosin were described previously.^18–20^ An enzyme system of pyruvate kinase and lactate dehydrogenase that regenerates ATP from the myosin-produced adenosine diphosphate (ADP) by oxidizing nicotinamide adenine dinucleotide (NADH) to the oxidized form of nicotinamide adenine dinucleotide (NAD). Conversion of NADH to NAD produces an absorbance change at 340 nm. The assay buffer consisted of PM12 buffer (12 mM PIPES (1,4-Piperazinediethanesulfonic acid), 2 mM magnesium chloride [MgCl_2_], 1 mM dithiothreitol, pH 6.8) supplemented with 60 mM potassium chloride (KCl) and an ATP concentration at ∼3-10× the K_M_ for the particular myofibril system (0.05 mM ATP for slow skeletal and cardiac, 0.5 mM ATP for fast skeletal). Separately, calcium concentrations were buffered using 0.6 mM Ethyleneglycol-*bis*(β-aminoethyl)-N,N,N□,N□-tetraacetic Acid (EGTA) and enough calcium chloride (CaCl_2_) to obtain the required free calcium concentration (calculated using a web resource: http://www.stanford.edu/~cpatton/webmaxc/webmaxcS.htm). Saturating concentration of blebbistatin was used to subtract non-myosin ATPase activity in cardiac and slow myofibrils. The concentration of 0.25 mg/mL was used for fast skeletal, whereas 1 mg/mL was used for cardiac and slow skeletal. Time-dependent absorbance measurements at 340 nm were carried out at ∼25 °C using either an Envision (Perkin-Elmer) or SpectraMax (Molecular Devices) plate reader. GraphPad Prism was used for data analysis.

Regulatory light chain (RLC) depletion in bovine cardiac myofibrils was performed as described previously.^22^ Myofibrils were treated with a buffer containing 20 mM Tris pH = 8.0, 10 mM trans-1,2-cyclohexanediaminetetraacetic acid (CDTA), 50 mM KCl, and 1% Trition X-100 for 30 minutes at room temperature. Subsequently, myofibrils were pelleted by centrifugation using an Eppendorf centrifuge 5417R and resuspended in the assay buffer described above for a washing step. Myofibrils were again centrifuged as before and resuspended in the assay buffer to perform an ATPase assay.

### Isothermal Titration Calorimetry Measurements

Myosin heavy chain fragments (HCFs) were expressed in Escherichia coli (E. coli) BL21 Star (Invitrogen) by glucose-lactose autoinduction^23^ using bicistronic expression vectors prepared by gene synthesis (ATUM.bio). RLC + 12 heptad-repeats (12-hep) HCF consisted of hMYL2 (UniProt P10916.3) expressed with an N-terminal maltose-binding protein (MBP) tag followed by a tobacco etch virus (TEV) protease cleavage site together with hMYH7 (UniProt P12883.5, K803-E927) expressed with an N-terminal 6×His tag followed by a TEV protease cleavage site. RLC + 2 heptad-repeats (2hep) HCF consisted of hMYL2 expressed with an N-terminal MBP tag followed by a TEV protease cleavage site together with hMYH7(K803-E855) expressed with an N-terminal 6×His tag followed by a TEV protease cleavage site. The RLC-only control consisted of hMYL2 expressed with an N-terminal 6×His and MBP tag followed by a TEV protease cleavage site. Proteins were purified by Ni-affinity chromatography (His-select resin, Sigma) followed by anion exchange (Source 15Q, Cytiva) in chromatography buffer (50 mM Tris-HCl, 2 mM MgCl_2_, 1 mM dithiothreitol [DTT], pH 8.0 at 4 °C), eluted with a gradient of sodium chloride (NaCl). Proteins were flash-frozen in liquid nitrogen following addition of 10% (w/v) sucrose.

Isothermal titration calorimetry (ITC) experiments were performed in a Micro-Cal AutoITC_200_ microcalorimeter (Malvern Panalytical, Inc.). Proteins were buffer exchanged into binding buffer (12 mM K-PIPES, 2 mM MgCl_2_, 60 mM KCl, 5 mM 2-mercaptoethanol, pH 6.8) by passage over a P-6DG column (Bio-Rad). Binding reactions were carried out in binding buffer containing 5 mM 2-mercaptoethanol and 3% dimethyl sulfoxide (DMSO). To correct for the heats of dilutions, the integrated heat signal from the last 2-3 injections were averaged and subtracted. Data collection and analysis was conducted using manufacturer-supplied Origin software using a single binding site model.

### Rat Cardiomyocyte Isolation, Contractility, and Calcium Transients

Rat cardiomyocytes were isolated from male ZSF1 lean and obese animals (25-30 weeks of age) using enzymatic digestion and Langendorff perfusion with a modified Krebs solution (perfusion buffer [PB]) containing (in mM): 10 dextrose, 135 NaCl, 4.7 KCl, 0.6 KH2PO4, 0.6 Na2HPO4, 1.2 MgSO4, 20 4-(2-Hydroxyethyl)piperazine-1-ethanesulfonic acid (HEPES), 30 taurine, and 10 2,3-butanedione monoxime (BDM) (pH 7.4). Briefly, the rats were heparinized (500 IU/mL) via intraperitoneal injection followed by anesthesia under 4% to 5% isoflurane. The heart was quickly excised from the anesthetized animal via lateral thoracotomy and the aorta cannulated. The coronary vessels were briefly flushed with perfusion buffer under retrograde perfusion at 37 °C followed by enzymatic digestion in PB containing 55 ug/mL Liberase TH (Roche) and 0.05 mM CaCl_2_. The isolated myocytes were collected in stop solution containing 50% fetal bovine serum (FBS) and allowed to settle by gravity. Calcium reintroduction was achieved through 3 subsequent washes with PB containing 0.5 mM, 0.75 mM, and 1.25 mM CaCl_2_, respectively. Cells were plated on laminin-coated (40 µg/mL) 35-mm glass-bottomed MatTek dishes in modified M199 medium containing 1× insulin transferrin selenium, 50 U/mL pen-strep, 5 mM taurine, 1 mM sodium pyruvate, 5 mM creatine, 2 mM carnitine, and 10 mM BDM. After a 2- to 3-hour recovery period, contractility and calcium transient measurements were made using an IonOptix Multicell system at 37 °C under continuous superfusion (∼1.5 mL/min) with pH 7.4 HEPES-buffered Tyrode’s solution containing (in mM): 140 NaCl, 5.4 KCl, 10 HEPES, 0.33 monosodium phosphate, 5 glucose, 1 MgCl_2_, and 1.8 CaCl_2_. For ratiometric calcium transient analysis, cells were loaded with 0.5 µM Fura2-AM (ThermoFisher) in Tyrode’s buffer for 13 minutes at room temperature. The cells were washed in Tyrode’s buffer and incubated at 37 °C for at least 20 minutes to ensure complete intracellular de-esterification. After achieving stable baseline measurements, ulacamten or DMSO vehicle controls were perfused, and subsequent repeated measures were obtained with increasing compound concentrations, after ∼5-minute equilibration between doses. All measurements were collected with 1 Hz field stimulation (20 V, 4 ms duration). Average transients were analyzed using CytoSolver software (IonOptix, version 2.0) to determine changes in contractility and/or calcium handling.

Normalized fractional shortening was calculated as: 100 × FS post-compound addition / FS pre-compound addition.

### Adult Human Cardiomyocyte Isolation and Contractility Analysis

Ulacamten dose response in human adult ventricular cardiomyocytes was performed by AnaBios Corporation (San Diego, CA). Apparent healthy normal donor hearts were obtained by legal consent and myocytes isolated using proprietary methods. Isolated cardiomyocytes were placed in a perfusion chamber mounted on an inverted Olympus IX83P1ZF microscope and allowed to equilibrate for 5 minutes under continuous superfusion (2 mL/min) with buffer temperature maintained at 35 ± 1 °C. After equilibration, myocytes were subjected to continuous field stimulation at 1 Hz frequency (3 ms duration, 1.5× voltage threshold required to initiate contraction). Sarcomere length transients were acquired with custom MyoBLAZER^TM^ acquisition software at an acquisition rate of 150 Hz using an Optromis Cyclone-25-150M camera. Repeated measures on selected cells were obtained with increasing concentrations of ulacamten to obtain dose-response curve, allowing 5-minute equilibration between doses. Optical density data corresponding to myocyte Z-lines was used to measure sarcomere length (SL) contractility via fast Fourier transform. To obtain an 8-point dose response, 2 separate batches of cells were evaluated at 4 overlapping doses for batch 1 (n = 6; 0.15, 0.6, 2.5, and 10 µM) and batch 2 (n = 7; 0.30, 1.25, 5, and 30 µM). Additional information on methods and experimental validation has been previously published.^24^

### Engineered Heart Tissues With Human Induced Pluripotent Stem Cell–Derived Cardiomyocytes

Dose-response analysis of ulacamten in human engineered heart tissues (EHTs) was performed by Propria LLC. Ventricular human induced pluripotent stem cell (hiPSC)-derived cardiomyocytes were differentiated from PGP1 wild-type hiPSCs or CRISPR-engineered PGP1 hiPSCs with heterozygous *MYH7 R403Q* mutation. EHTs were formed in custom MyoPod scaffolds by seeding hiPSC-derived cardiomyocytes and primary human cardiac fibroblasts (hCFs) in decellularized porcine myocardial scaffolds at a ratio of 1:10 (1 × 10^6^ hiPSC cardiomyocytes and 10 × 10^4^ hCFs per tissue). Tissues were cultured for 2-3 weeks, followed by 6-point compound dose-response testing in Propria’s MyoLab automated contractility device. All measurements were performed at physiologic temperature (37 °C) with 1 Hz frequency stimulation and media superfusion (Dulbecco’s Modified Eagle Medium (DMEM) + 0.01% DMSO) at 0.3 mL/min. Three 5- to 10-second recordings were collected in each condition, and custom analysis software was used to calculate force (µN), time to peak force (TTP [ms]), and return time (ms) to 50% and 90% (RT_50_ and RT_90_).

### Effect of Ulacamten in ZSF1 Rat Model

Eight-week-old male ZSF1 lean and obese rats were obtained from Charles River Laboratories. At 20 weeks of age, ZSF1 obese rats received either vehicle (0.5% hydroxypropyl methylcellulose (HPMC)/0.1% Tween suspension mixed with peanut butter) or ulacamten (10 mg/kg in 0.5% HPMC/0.1% Tween suspension mixed with peanut butter) daily for 12-14 weeks. From arrival to the end of study, echocardiograms were performed to assess heart rate, cardiac dimensions, and measures of systolic and diastolic function every 4 weeks. At 32-34 weeks of age, all animals were euthanized. Hearts were weighed, fixed in 10% formalin, and embedded in paraffin. The 10 µm–thick cross-sections from the mid-ventricle region were stained for picrosirius red and Masson’s trichrome stain. Images were acquired and analyzed by a Keyence BZ-X and accompanying software.

### Statistical Analyses

All data are presented as mean ± standard deviation (SD) or mean ± standard error (SE), with statistical analyses performed using GraphPad Prism software. Sample sizes and individual statistical tests used for each experiment are detailed in the relevant figure legends.

## RESULTS

### Ulacamten Partially Inhibits the ATPase Activity of Myosin

The precursor of ulacamten was discovered in a high-throughput biochemical screen such as previously reported for aficamten.^18^ Optimization of the chemical structure for desirable drug-like properties led to the discovery of ulacamten (Supplemental Figure 1). A distinguishing feature of this class of CMIs was the partial inhibition of myofibrillar ATPase activity (∼50%) observed at concentrations >10 µM (**Figures 1A and 1B**). This property made ulacamten distinct in comparison to aficamten and mavacamten, which have complete^18^ or nearly complete^25^ maximal inhibition of cardiac myofibril ATPase activity. The inhibition of the myofibrillar ATPase by ulacamten was stereoselective in that CK-4022425, a diastereomer of ulacamten, did not inhibit the myofibrillar ATPase (Supplemental Figure 1), indicating binding of ulacamten was dependent on the specific conformation of ulacamten. The biochemical EC_50_ (concentration at which inhibition is 50% of maximal activity) of ulacamten in assays using bovine cardiac myofibrils was 2.9 μM (95% CI: 2.74-3.16) (**Figure 1A**, Supplemental Table 1) as compared with 11.9 μM (95% CI: 9.3-17.8) with rabbit fast skeletal myofibrils (Supplemental Figure 2A, Supplemental Table 1), indicating selectivity for cardiac vs fast skeletal muscle types. The EC_50_ in assays using bovine slow skeletal myofibrils was 2.5 μM (95% CI: 1.65-4.64) (Supplemental Figure 2A, Supplemental Table 1), similar to cardiac myofibrils, as would be expected for a myosin-targeted inhibitor because cardiac and slow skeletal muscles express the same myosin isoform.^26^

**Figure 1.**
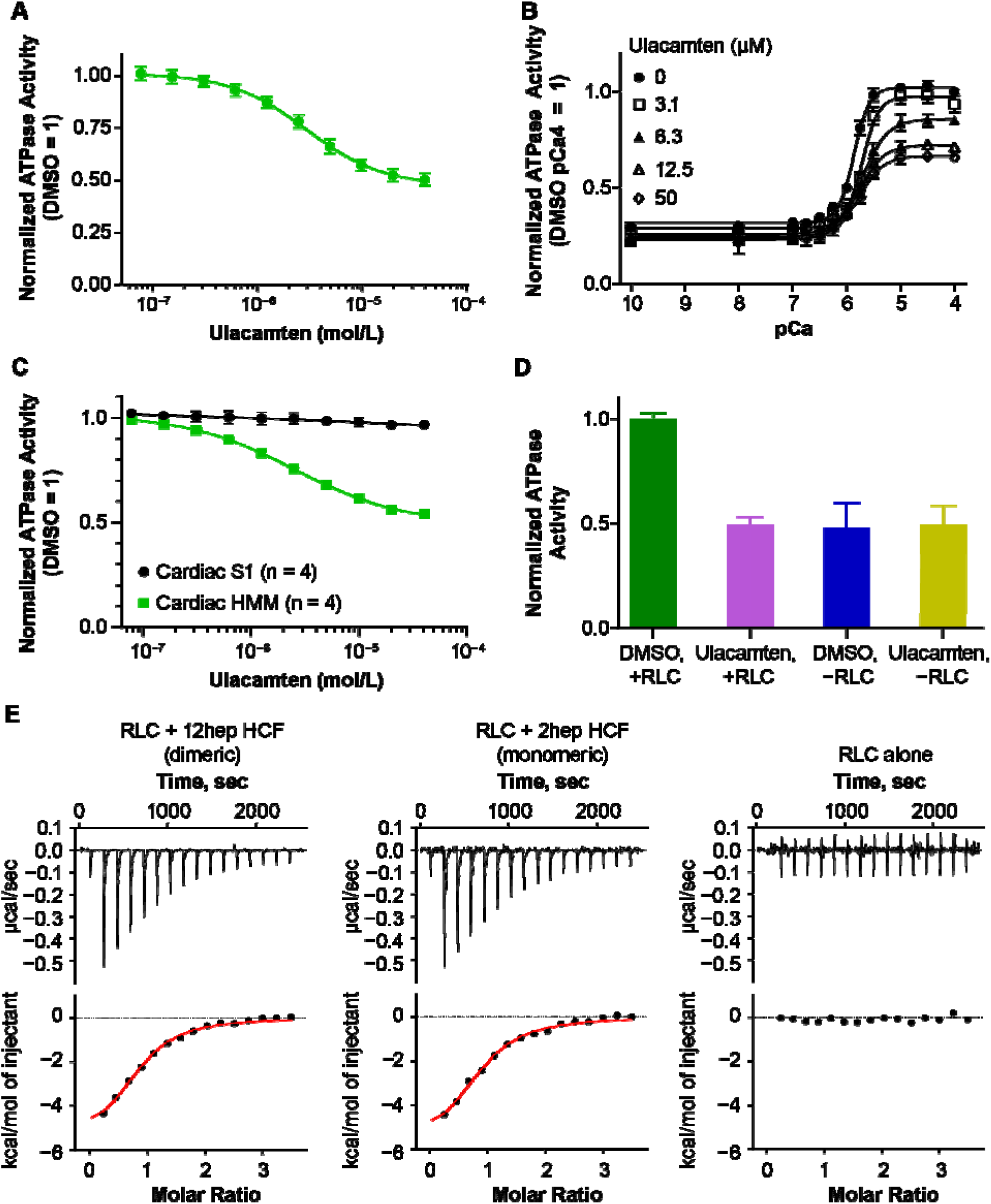
Biochemical and Biophysical Characterizations of Cardiac Myosin Binding to Ulacamten. (A) Inhibition of myosin ATPase activity in bovine cardiac myofibrils by ulacamten. ATPase activities were measured at the calcium ion concentration of 75% activation. Data represent mean ± SD with four-parameter fitting. The biochemical EC_50_, which represents the ulacamten concentration for 50% of the maximal inhibition, was 2.9 μM (n = 31, 95% CI 2.74-3.16). (B) Calcium dependence of bovine cardiac myofibril ATPase inhibition. Data shown are mean ± SD with four-parameter fitting. (C) Comparison of myosin inhibition in single-headed chymotryptic subfragment-1 (S1) and two headed heavy meromyosin (HMM) by ulacamten. Dose-response curves of actin-activated myosin ATPase activity are shown. Data represent mean ± SD with four-parameter fittings. (D) Testing the presence of regulatory light chain (RLC) for the myosin ATPase inhibition by ulacamten in bovine cardiac myofibrils. Myofibrils were treated with Triton X-100 for the depletion of RLC (see Methods); 1 mM calcium chloride (CaCl_2_) was added in myofibrils before ATPase activity measurements. ATPase activities are shown in the presence of 1% Dimethyl Sulfoxide (DMSO) or 50 μM ulacamten in both intact myofibrils ([DMSO: +RLC (n = 24); ulacamten: +RLC (n = 31)]) and RLC-depleted myofibrils ([DMSO: −RLC (n = 31); ulacamten: −RLC (n = 32)]). Raw ATPase rates were normalized to reactions containing an equivalent concentration of DMSO in myofibrils with RLC. (E) Binding of ulacamten to recombinant myosin heavy chain fragments (HCF) measured by isothermal titration calorimetry (ITC). Data shown are a single representative binding isotherm. Aggregate data are presented in Table S1.

ATPase assays using purified actin and myosin were employed to identify the target of ulacamten and to explore its mechanism of action. Existing CMIs are complete inhibitors of the ATPase of the single-headed S1 fragment ATPase of myosin. Surprisingly, ulacamten failed to show any effects in single-headed chymotryptic S1 (**Figure 1C**) and papain S1 (Supplemental Figure 2B) at concentrations as high as 40 μM, suggesting a requirement for an intact 2-headed myosin to exert its effect. The EC_50_ in ATPase assays using 2-headed bovine cardiac HMM was 2.5 μM (95% CI: 2.3-2.8) with a maximal inhibition of ∼50%, both similar to what was observed in bovine cardiac myofibrils (**Figure 1C**, Supplemental Table 2). This supports myosin as the target of ulacamten and suggests the partial inhibition observed could be due to a unique conformation of myosin and not simply an artifact of the myofibril system. Inhibition of smooth muscle myosin is undesirable given the potential effect to lower blood pressure. Ulacamten does not inhibit the actin-activated ATPase of chicken gizzard smooth muscle HMM at concentrations up to 40 μM (Supplemental Figure 2C), demonstrating selectivity for cardiac myosin.

### The RLC Region of Myosin Is the Binding Site of Ulacamten

Biochemical and biophysical experiments were performed to understand the binding site of ulacamten. The ATPase inhibition observed in 2-headed HMM but not in single-headed S1 (**Figure 1C**) suggested the ulacamten binding site region might extend beyond the motor domain responsible for binding to actin and ATP.^27^ The motor domain is connected to the proximal tail region though the lever arm region where the essential light chain (ELC) and regulatory light chain (RLC) bind. We first tested whether RLC was required for ulacamten binding. Myofibrils were treated with Triton X-100 and CDTA^22^ to deplete the RLC, which was confirmed by SDS PAGE (Sodium Dodecyl Sulfate–Polyacrylamide Gel Electrophoresis) (Supplemental Figure 2D). RLC depletion resulted in a partial loss of myosin ATPase activity (**Figure 1D**), consistent with previous literature.^22^ Interestingly, ulacamten treatment of RLC-depleted myofibrils did not further inhibit myosin ATPase activity, indicating the presence of RLC was required for ulacamten activity and suggesting the RLC may comprise at least part of the ulacamten binding site (**Figure 1D**).

The binding site was further defined using ITC to measure the thermodynamics of ulacamten binding to recombinant cardiac myosin heavy chain fragments (HCFs), based on previous studies used to describe the binding site for piperine on fast skeletal myosin.^28^ These myosin fragments were produced in E. coli by co-expression of the human RLC and a portion of human β-cardiac myosin that lacks the motor domain but includes the RLC-binding region and varying lengths of the coiled-coil proximal tail. A longer length of tail comprising 12 heptad repeats (12-hep HCF) dimerizes and models the lever arm of 2-headed HMM, whereas a shorter tail length comprising two heptad repeats (2hep HCF) models single-headed S1. Binding of ulacamten to either 12-hep HCF or 2hep HCF elicited a clear exothermic signal of 6.1 kcal/mol, and fitting of the binding data with a single-site model indicated an affinity of 5.3 µM and a stoichiometry consistent with a single binding on each RLC-containing monomer (**Figure 1E**, Supplemental Table 3). No significant binding was observed when the RLC was tested in the absence of the myosin HCF (**Figure 1E**)..

The ability of ulacamten to bind to 2hep HCF was surprising, given the lack of ATPase inhibition observed with S1. To confirm this finding, binding experiments were performed with bovine cardiac papain S1 containing an intact RLC.^29^ Clear exothermic heat signals of 5.7 kcal/mol were observed, consistent with the binding heats observed for RLC + 2hep HCF (Supplemental Figure 3A, Supplemental Table 4). The binding affinity of ulacamten for papain S1 (3.9 µM; Supplemental Table 4) was similar to the ATPase EC_50_ in myofibrils (2.9 µM), suggesting that a binding reaction relevant to functional modulation was observed. As a control, binding experiments were performed with an inactive isomer of ulacamten (CK-4022425; Supplemental Figure 1). No significant binding was observed between 12-hep HCF, 2hep HCF, or RLC proteins and the inactive isomer (Supplemental Figure 3B). Taken together with the RLC-depletion experiments above, these data suggest that although single-headed myosin can bind ulacamten, inhibition of ATPase activity requires the presence of both the RLC and 2 myosin heads.

### Ulacamten Inhibits Cellular Contractility Across Multiple Species and Disease States

To evaluate cellular potency of ulacamten, we measured sarcomere length shortening and calcium transients in isolated myocytes from the ZSF1 obese rats. Obese ZSF1 rats are a model of HFpEF with diastolic dysfunction, elevated N-terminal pro-B-type natriuretic peptide (NT-proBNP), cardiac hypertrophy, and inflammation potentially secondary to the comorbidities of obesity, hyperglycemia, and hypertension.^30^ Myocytes isolated from ZSF1 obese rats exhibit impaired contraction and relaxation kinetics^31^ and dysfunctional calcium regulation.^32^ Ulacamten demonstrated potent inhibition of sarcomere length shortening in myocytes from both ZSF1 obese rats and ZSF1 lean controls, without affecting calcium transients (**Figures 2A and 2B**, Supplemental Figure 4). The cellular potency was consistent with biochemical EC_50_ values and comparable between ZSF1 phenotypes, with half-maximal inhibitory concentration (IC_50_) values of 3.2 µM in obese and 2.6 µM in lean myocytes (**Figure 2B**, Supplemental Figure 4B). Similar potency was observed in adult human myocytes, with an IC_50_ value of 2.9 µM (**Figure 2C**). To evaluate ulacamten in the context of human disease with hypercontractility, EHTs were produced using laser-cut decellularized porcine myocardium scaffolds and iPSC-cardiomyocytes carrying the HCM-associated R403Q myosin mutation or isogenic controls (wild-type). Introduction of R403Q mutation resulted in an alteration in contractile parameters consistent with hypercontractile system with impaired relaxation parameters (Supplemental Figure 5). In both wild-type and R403Q EHTs, treatment with ulacamten resulted in >80% force inhibition at 10 µM, with IC_50_ values of 0.85 µM and 3.08 µM, respectively (**Figure 2D**). Ulacamten inhibited all the contractile properties in a dose-dependent manner (Supplemental Figure 6). In R403Q EHTs, at concentrations approximately the IC_50_ value (∼2 µM), ulacamten restored contractile force and relaxation times (both RT_50_ and RT_90_) in R403Q EHTs to near wild-type levels (**Figure 2E**, Supplemental Figure 7).

**Figure 2.**
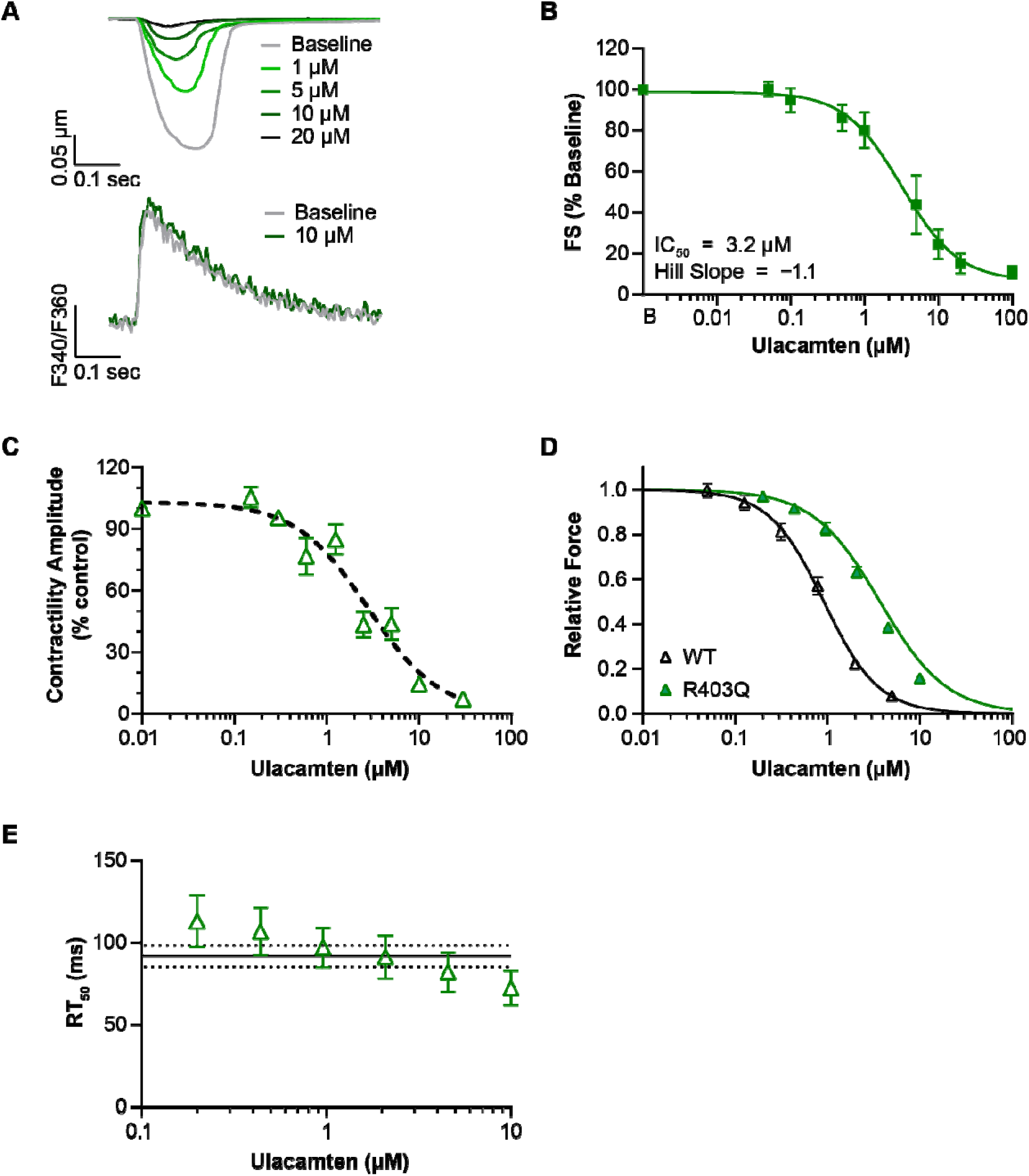
Functional Characterization of Ulacamten on Cardiomyocyte Contractility. (A) Representative traces of sarcomere length (SL) shortening (top) and ratiometric Fura-2 calcium transients (bottom) in ZSF1 obese rat myocytes treated with escalating concentrations of ulacamten. (B–C) Ulacamten dose-response inhibition of SL fractional shortening (FS) in (B) rat ZSF1 obese (n = 4 isolations, >40 cells) and (C) isolated adult human ventricular myocytes (n = 13 cells from 2 donors). Data are normalized to baseline FS prior to compound addition and fit with 4-parameter dose-response curves to determine 50% of the maximal inhibition (IC_50_) and Hill slope. (D–E) Ulacamten dose-response inhibition of engineered heart tissue (EHTs) derived from wild-type (WT) (n = 5) and R403Q iPSC cardiomyocytes (n = 10). (D) Relative force normalized to baseline values. (E) Return time to 50% baseline (RT_50_; ms) for R403Q EHTs. Solid and dotted lines represent the value for wild-type and its standard deviation, respectively.

### Ulacamten Reduced Fractional Shortening and Improved Relaxation in ZSF1 Obese Rats

The effect of chronic ulacamten treatment was evaluated in the ZSF1 obese HFpEF rat model. At 20 weeks of age, ZSF1 obese rats received either vehicle or ulacamten (10 mg/kg by mouth [PO] once daily [QD]) for 12-14 weeks. At the time of treatment initiation, ZSF1 obese rats had significantly lower heart rate, greater LV wall thickness, and impaired diastolic function (**Table 1**). Relative to vehicle-treated ZSF1 obese rats, treatment for 12+ weeks of daily ulacamten significantly reduced cardiac fractional shortening (vehicle 51.6 ± 1.8% vs ulacamten 44.0 ± 1.6%, *P* < 0.0001; **Figure 3A**). In association with lower fractional shortening, reductions in wall thickness (vehicle 1.88 ± 0.1 mm vs ulacamten 1.59 ± 0.1 mm, *P* ≤ 0.0001 for posterior wall thickness at diastole, vehicle 1.94 ± 0.1 mm vs ulacamten 1.60 ± 0.1 mm, *P* ≤ 0.0001 for anterior wall thickness at diastole; **Table 2**) and isovolumic relaxation time (IVRT; vehicle 27.8 ± 1.7 ms vs ulacamten 25.2 ± 1.6 ms, *P* < 0.01; **Figure 3B**, **Table 2**) were observed with ulacamten treatment. At the end of the study, hearts were collected for histologic analysis of fibrosis. Picrosirius red stains of heart cross-sections revealed significant collagen content in vehicle-treated ZSF1 obese rats compared with ZSF1 lean rats (obese 5.5 ± 0.6% vs lean 3.2 ± 0.6%, *P* < 0.0001; **Figures 3C and 3D**, **Table 2**). Cardiac tissue from ulacamten-treated ZSF1 obese rats demonstrated significantly lower collagen staining relative to vehicle-treated ZSF1 obese rats (ulacamten 3.2 ± 0.4% vs vehicle 5.5 ± 0.6%, *P* < 0.0001; **Figures 3C and 3D**, **Table 2**).

**Figure 3.**
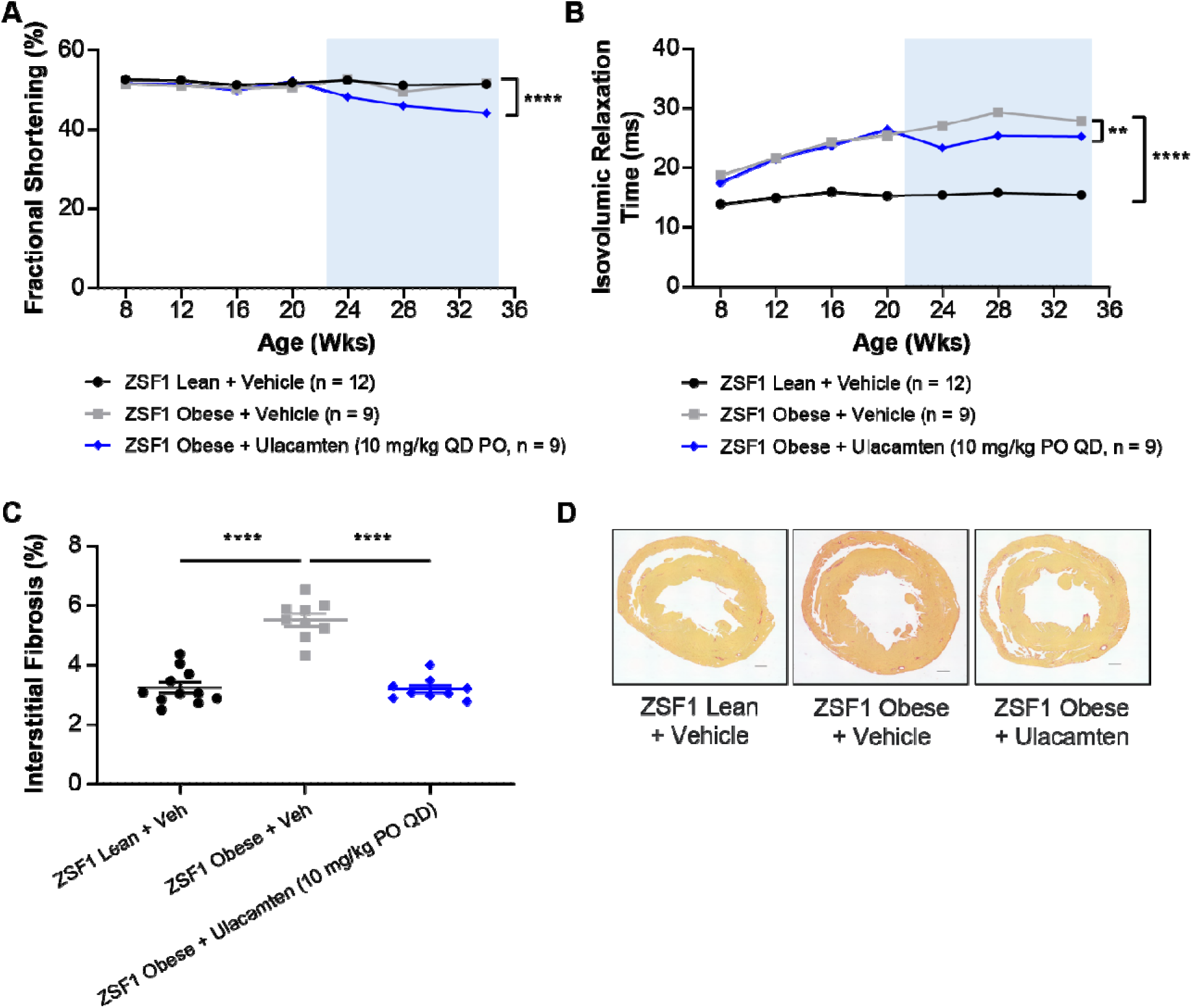
Chronic Ulacamten Treatment Improves Cardiac Relaxation in ZSF Obese Rats. (A) Ulacamten treatment (10 mg/kg orally once daily [QD PO]) reduced rat cardiac fractional shortening in ZSF1 obese rats. (B) Isovolumic relaxation time (IVRT) is lower in ZSF1 obese rats treated with ulacamten. (C) ZSF1 obese rats dosed with ulacamten (10 mg/kg, QD PO) for 12-14 weeks demonstrate lower interstitial fibrosis than vehicle-treated ZSF1 obese rats. (D) Representative picrosirius red-stained heart sections for collagen deposition from ZSF1 Lean and Obese rats with and without ulacamten treatment. All data is expressed as mean ± standard error of the mean. \*\**P* < 0.001, \*\*\*\**P* < 0.0001.

**TABLE 1.**
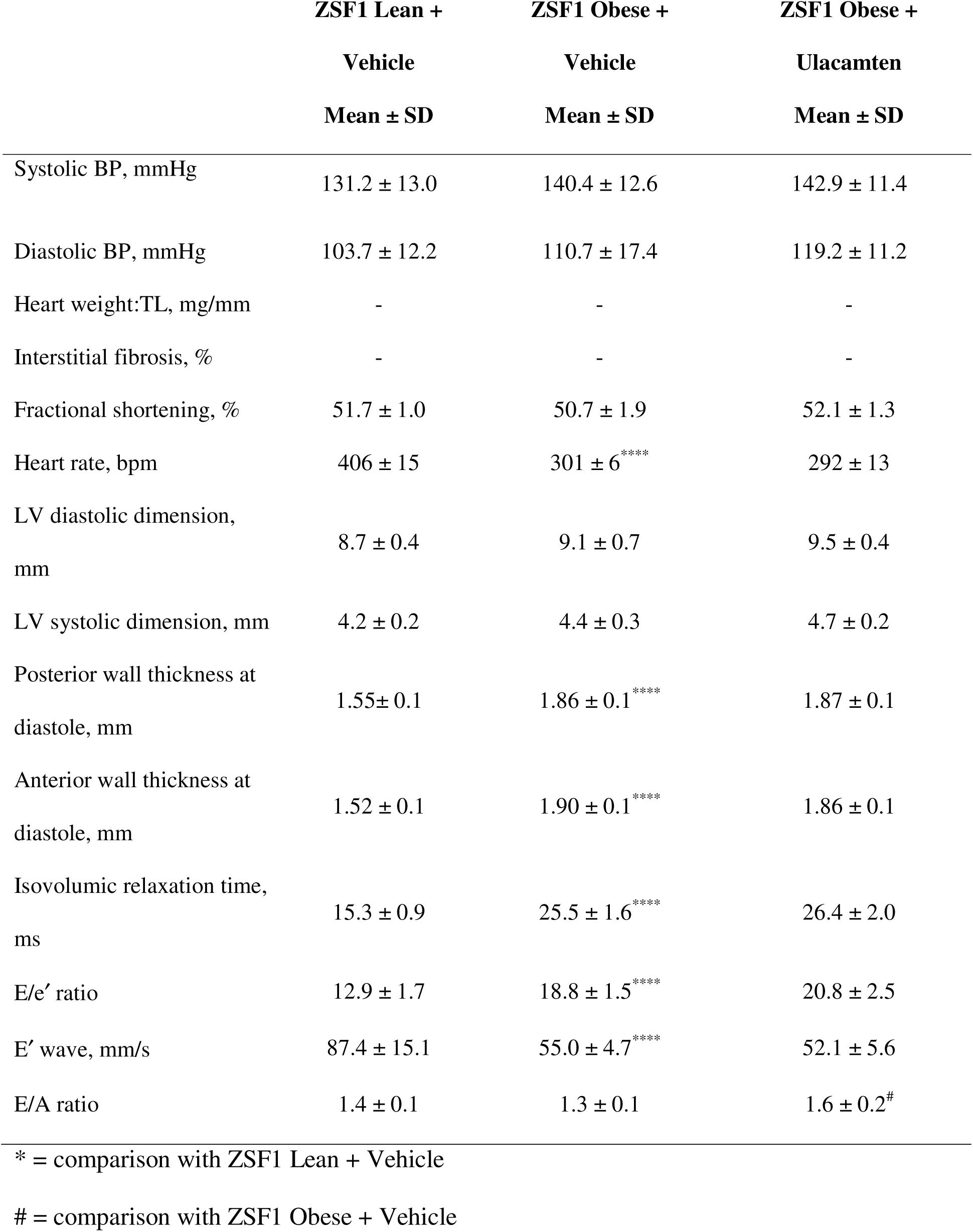
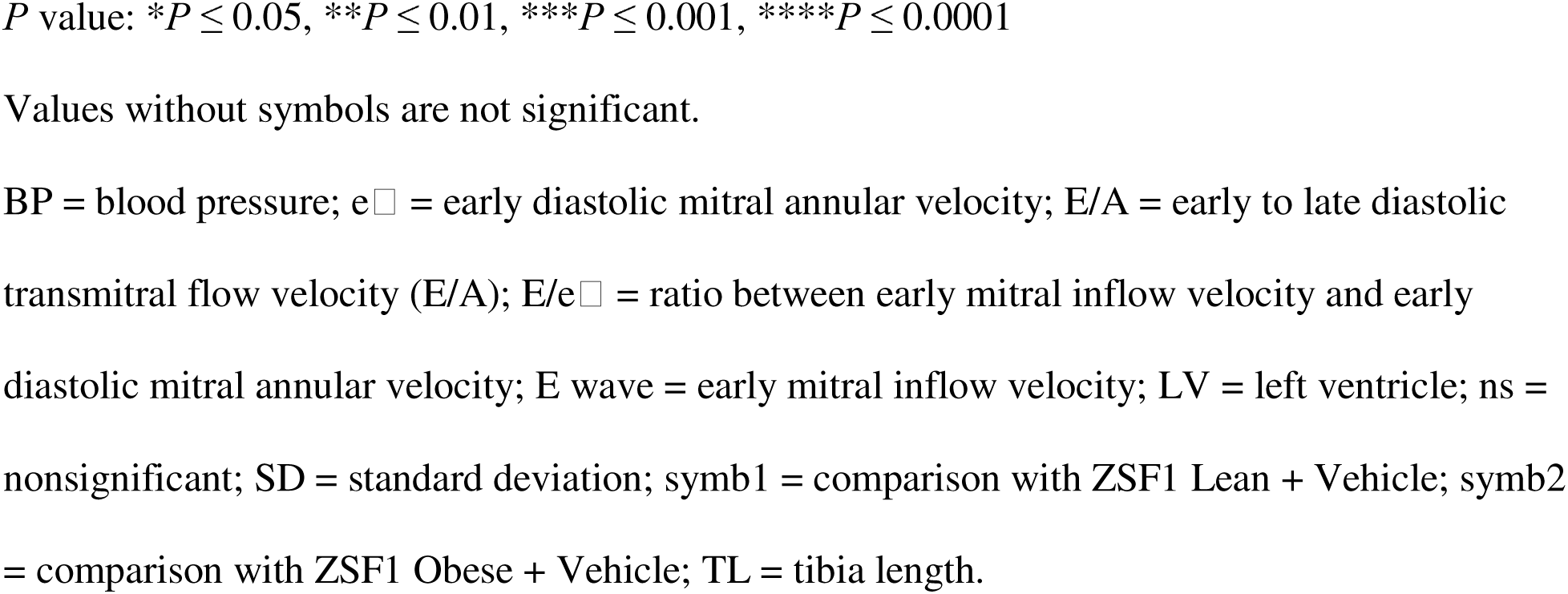
Baseline Characteristics of ZSF1 Rats at 20 Weeks.

|  | **ZSF1 Lean +**<br>**Vehicle**<br>**Mean ± SD** | **ZSF1 Obese +**<br>**Vehicle**<br>**Mean ± SD** | **ZSF1 Obese +**<br>**Ulacamten**<br>**Mean ± SD** |
| --- | --- | --- | --- |
| Systolic BP, mmHg | 131.2 ± 13.0 | 140.4 ± 12.6 | 142.9 ± 11.4 |
| Diastolic BP, mmHg | 103.7 ± 12.2 | 110.7 ± 17.4 | 119.2 ± 11.2 |
| Heart weight:TL, mg/mm | - | - | - |
| Interstitial fibrosis, % | - | - | - |
| Fractional shortening, % | 51.7 ± 1.0 | 50.7 ± 1.9 | 52.1 ± 1.3 |
| Heart rate, bpm | 406 ± 15 | 301 ± 6<sup>****</sup> | 292 ± 13 |
| LV diastolic dimension,<br>mm | 8.7 ± 0.4 | 9.1 ± 0.7 | 9.5 ± 0.4 |
| LV systolic dimension, mm | 4.2 ± 0.2 | 4.4 ± 0.3 | 4.7 ± 0.2 |
| Posterior wall thickness at<br>diastole, mm | 1.55 ± 0.1 | 1.86 ± 0.1<sup>****</sup> | 1.87 ± 0.1 |
| Anterior wall thickness at<br>diastole, mm | 1.52 ± 0.1 | 1.90 ± 0.1<sup>****</sup> | 1.86 ± 0.1 |
| Isovolumic relaxation time,<br>ms | 15.3 ± 0.9 | 25.5 ± 1.6<sup>****</sup> | 26.4 ± 2.0 |
| E/e' ratio | 12.9 ± 1.7 | 18.8 ± 1.5<sup>****</sup> | 20.8 ± 2.5 |
| E' wave, mm/s | 87.4 ± 15.1 | 55.0 ± 4.7<sup>****</sup> | 52.1 ± 5.6 |
| E/A ratio | 1.4 ± 0.1 | 1.3 ± 0.1 | 1.6 ± 0.2<sup>#</sup> |

\* = comparison with ZSF1 Lean + Vehicle

**TABLE 2.**
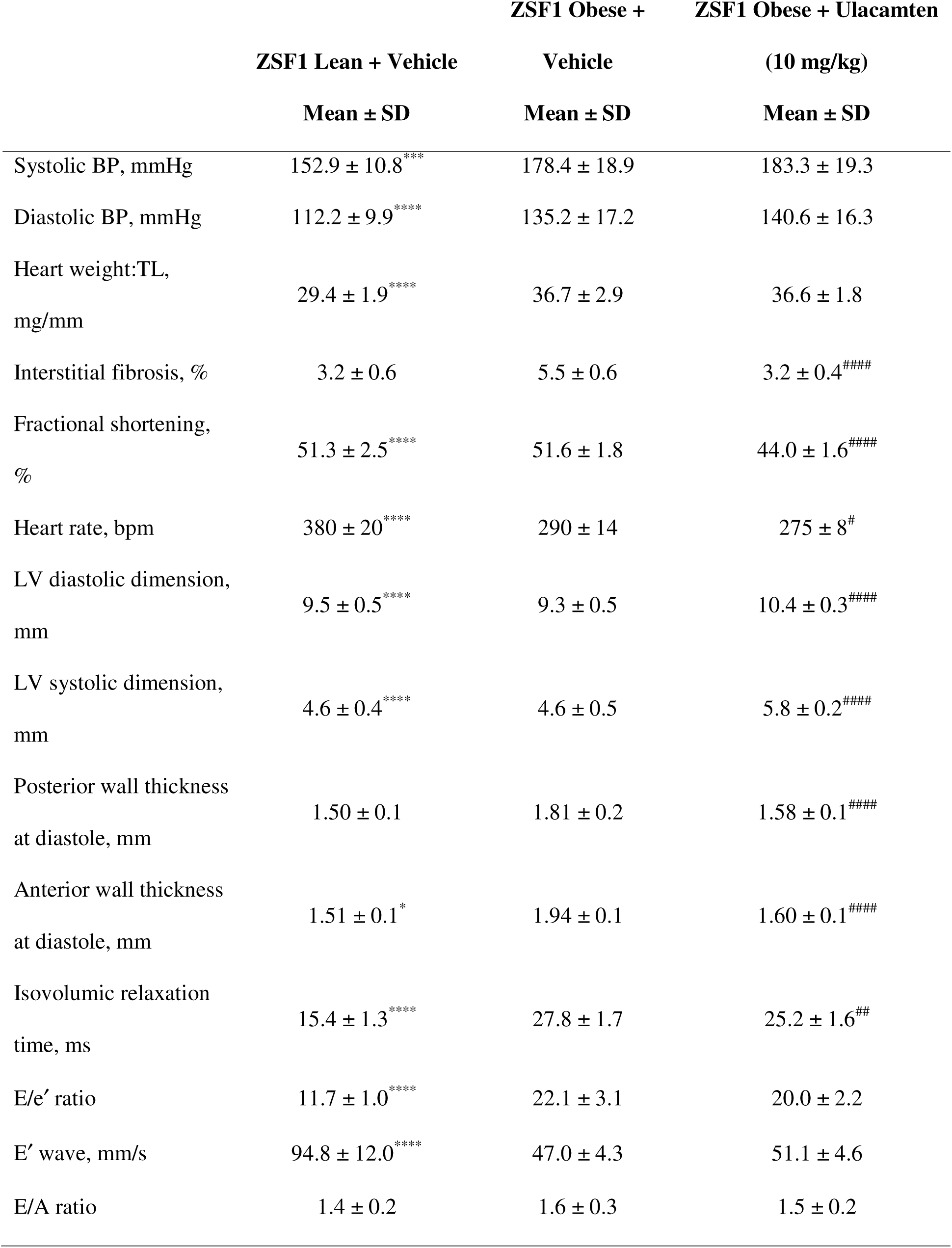
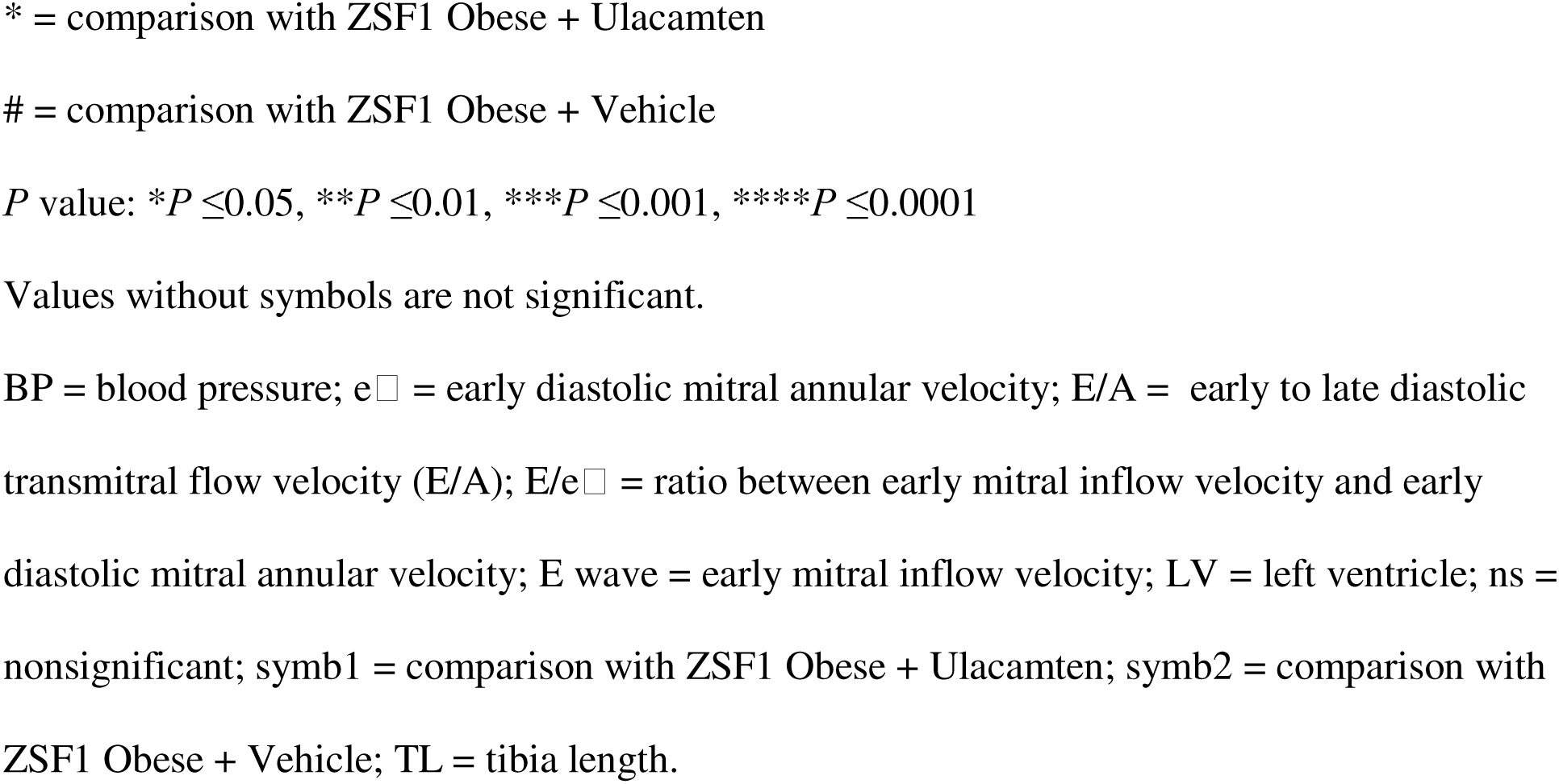
Endpoint Characteristics of ZSF1 Rats at 34 Weeks.

|  | **ZSF1 Lean + Vehicle** | **ZSF1 Obese +**<br>**Vehicle** | **ZSF1 Obese + Ulacamten**<br>**(10 mg/kg)** |
| --- | --- | --- | --- |
|  | **Mean ± SD** | **Mean ± SD** | **Mean ± SD** |
| Systolic BP, mmHg | 152.9 ± 10.8<sup>***</sup> | 178.4 ± 18.9 | 183.3 ± 19.3 |
| Diastolic BP, mmHg | 112.2 ± 9.9<sup>****</sup> | 135.2 ± 17.2 | 140.6 ± 16.3 |
| Heart weight:TL,<br>mg/mm | 29.4 ± 1.9<sup>****</sup> | 36.7 ± 2.9 | 36.6 ± 1.8 |
| Interstitial fibrosis, % | 3.2 ± 0.6 | 5.5 ± 0.6 | 3.2 ± 0.4<sup>####</sup> |
| Fractional shortening,<br>% | 51.3 ± 2.5<sup>****</sup> | 51.6 ± 1.8 | 44.0 ± 1.6<sup>####</sup> |
| Heart rate, bpm | 380 ± 20<sup>****</sup> | 290 ± 14 | 275 ± 8<sup>#</sup> |
| LV diastolic dimension,<br>mm | 9.5 ± 0.5<sup>****</sup> | 9.3 ± 0.5 | 10.4 ± 0.3<sup>####</sup> |
| LV systolic dimension,<br>mm | 4.6 ± 0.4<sup>****</sup> | 4.6 ± 0.5 | 5.8 ± 0.2<sup>####</sup> |
| Posterior wall thickness<br>at diastole, mm | 1.50 ± 0.1 | 1.81 ± 0.2 | 1.58 ± 0.1<sup>####</sup> |
| Anterior wall thickness<br>at diastole, mm | 1.51 ± 0.1<sup>*</sup> | 1.94 ± 0.1 | 1.60 ± 0.1<sup>####</sup> |
| Isovolumic relaxation<br>time, ms | 15.4 ± 1.3<sup>****</sup> | 27.8 ± 1.7 | 25.2 ± 1.6<sup>##</sup> |
| E/e' ratio | 11.7 ± 1.0<sup>****</sup> | 22.1 ± 3.1 | 20.0 ± 2.2 |
| E' wave, mm/s | 94.8 ± 12.0<sup>****</sup> | 47.0 ± 4.3 | 51.1 ± 4.6 |
| E/A ratio | 1.4 ± 0.2 | 1.6 ± 0.3 | 1.5 ± 0.2 |

\* = comparison with ZSF1 Obese + Ulacamten

## DISCUSSION

Successful clinical trials of mavacamten and aficamten in oHCM have demonstrated the clinical utility of CMIs, igniting interest in discovering and developing additional CMIs for patient populations beyond oHCM. We have shown that ulacamten is a new class of cardiac myosin inhibitor that has a mechanism of action distinct from both mavacamten and aficamten.

Aficamten binds to myosin between the U50 (upper 50 kDa) and the L50 (lower 50 kDa) subdomains, near the inorganic phosphate (Pi)-release backdoor.^18^ Aficamten binding stabilizes a pre-power stroke conformation that hinders myosin heads entering the mechanochemical cycle of force production. Mavacamten binds to a different site of the motor domain, between the N-terminal (N-term) and the Converter subdomains of the motor domain, stabilizing a pre-power stroke conformation that promotes the interacting-heads motif (IHM) geometry of 2-headed myosin where the 2 heads bind to its own proximal tail to form a folded state.^33–36^ Ulacamten, on the contrary, binds outside the catalytic domain of the myosin head, requiring both the regulatory light chains and the RLC-binding region of the myosin heavy chain. Thus, surprisingly, there appear to be 3 distinct binding sites for small molecules that can inhibit the cardiac myosin ATPase.

Despite the differences among the binding sites of CMIs, all are inhibitors of contractile force in cardiomyocytes^18^ or fibers.^18,37^ Ulacamten inhibits fractional shortening to near completion at concentrations >10 μM, in rat and human primary cardiomyocytes as well as EHT created with human iPSC-derived cardiomyocytes. As CMIs reduce the number of heads undergoing cross-bridge formation, there is a loss of motion between the thick and thin filaments, even when the latter are activated by calcium. For mavacamten and aficamten, a reduction in the number of active cross-bridges is reflected in the reduction of the ATPase rate in myofibrillar assays. Further, with these compounds, the concentration dependence of myofibrillar ATPase inhibition closely follows that of cellular contractility. The inhibition of myosin ATPase was almost complete for mavacamten and aficamten in both myofibrillar and purified myosin at concentrations >10 μM.^18,25^ In contrast, ulacamten demonstrates only partial inhibition at saturating doses in cardiac myofibrils, which has also been observed with another recently reported CMI.^38^ Two-headed HMM also retains the partial biochemical inhibition of the actin-activated ATPase activity at saturating concentrations of ulacamten. Together, these observations indicate that the formation of a partially inhibited state by ulacamten in myofibrils is independent of other sarcomeric proteins.

Ulacamten binds to single-headed S1 but cannot inhibit its ATPase activity. Interestingly, ulacamten binds to RLC in complex with short myosin HCFs, but not to RLC alone. Taken together with the RLC-depletion experiments, this suggests RLC is necessary but not sufficient for ulacamten binding and that binding to single-headed myosin is not sufficient for functional inhibition of ATPase activity. For mavacamten, X-ray fiber diffraction studies indicate that mavacamten binding leads to the folding of myosin back into the thick filaments,^35^ and this conformational change is associated with loss of ATPase activity. In contrast, similar studies with aficamten indicate that the position of aficamten-bound myosin heads remain unbound to the thick filament,^39^ and instead, aficamten stabilizes a unique head conformation of myosin that prevents entry into the mechanochemical cycle and binding to actin without docking the heads on the thick filament of myosin. A detailed structural study of cardiac fibers with myosin bound to ulacamten will help to further elaborate its mechanism of inhibition.

The efficacy of ulacamten was tested in R403Q EHTs as a hypercontractile in vitro cardiac tissue model. Importantly, R403Q EHTs demonstrated slower relaxation kinetics, as manifested in an increase in both RT_50_ and RT_90_ values with respect to wild-type EHTs. Both wild-type and R403Q EHTs showed dose-response curves of force consistent with those measured in primary cardiomyocytes from humans and from the ZSF1 rat model with similar IC_50_ values. Treating R403Q EHTs with ulacamten improved both RT_50_ and RT_90_ even at concentrations where both force and time to peak force have minimal alterations.

The ZSF1 rat model recapitulates aspects of HFpEF with comorbidities, including hypertension, diabetes, and obesity, that contribute to a phenotype that includes LV hypertrophy, diastolic dysfunction, and fibrosis. Ex vivo muscle mechanics measurements in permeabilized ZSF1 rat cardiomyocytes showed elevated passive tension that was similar to HFpEF patient-derived tissue samples, owing to excessive cross-bridge fomation.^40^ Previous reports have demonstrated efficacy in the ZSF1 obese model with clinically validated therapeutics.^41,42^ In this current study, ZSF1 obese rats demonstrated increased wall thickness, IVRT, and E/e’ at 20 weeks of age in comparison to ZSF lean rat controls. Following 12-14 weeks of ulacamten treatment (10 mg/kg PO QD), a decrease in fractional shortening by ulacamten was correlated with improvements in LV dimensions, wall thickness, IVRT, and interstitial fibrosis with respect to age-matched, vehicle-treated ZSF1 obese rats. The improvement in function by ulacamten parallels the reported improvement by the cardiac myosin inhibitor, mavacamten, in the same rat model and in HFpEF patient biopsies.^40^

The complexity in the pathophysiological changes of the heart in HFpEF^43,44^ might have complicated the development of drugs for the entire range of EF observed in patients with HFpEF. Many heart failure clinical trials included patients with HFmrEF and HFpEF, relying on the post hoc analysis of trials for the extraction of benefits for these 2 individual subgroups.

SGLT2 inhibitors are the only class of drugs suitable for all patients with HFpEF, whereas angiotensin receptor-neprilysin inhibitors and mineralocorticoid receptor antagonists are restricted for certain patients.^45^ Recent clinical agents like glucagon-like peptide-1 (GLP-1) agonists have shown benefits in cardiometabolic outcomes in HFpEF clinical trials, but their efficacy in non-obese patients will require further testing.^46^ The HFpEF clinical trial (NCT06215586) of the phosphodiesterase 9 inhibitor CRD-750 excludes the patient population with EF >60%. Mavacamten treatment has shown improvements in cardiac biomarkers as well as NYHA functional class among patients with HFpEF and EF >60% in an open-label, single-arm phase 2 clinical trial (EMBARK-HFpEF, NCT04766892).^47^ Ulacamten is currently being tested in a double-blinded, randomized clinical trial among patients with symptomatic heart failure and LVEF ≥60% (AMBER-HFpEF, NCT06793371).

There are a few limitations in this study. First, we designed biochemical and biophysical measurements to understand the binding site of ulacamten in myosin. A complete understanding, however, requires further work with structural tools like X-ray fiber diffraction or Cryo-EM to elucidate the mechanistic details of myosin inhibition by ulacamten. Second, we studied improvements in relaxation parameters by the action of ulacamten in R403Q EHTs although the patient population in HFpEF clinical trials is not expected to carry HCM mutations. This model was used here to demonstrate the relative dependence of RT_50_ or RT_90_ on ulacamten concentration with respect to the decline of force, which can be an indicator of systolic function in vivo. Of note, the R403Q mutation lies outside the RLC-binding region, and the comparison between wild-type and R403Q EHTs should not be complicated by the alteration of ulacamten affinity due to the change in amino acid sequence. Third, we tested ulacamten treatment in ZSF1 rats because it has been widely used in preclinical testing other agents in development for treating HFpEF. Given the complexity of HFpEF pathophysiology, this rat model recapitulates some of the phenotype of HFpEF but not all. The ongoing clinical trial of ulacamten will ultimately validate the translational value of our ZSF1 rat study.

## CONCLUSIONS

Our study delineates the mechanism of action of ulacamten and establishes it as a distinct myosin inhibitor in comparison to mavacamten and aficamten. Ulacamten binds to the RLC-binding region of myosin, leading to an inhibitory state of myosin that has lost ∼50% of ATPase activity in both myofibrillar and purified protein preparations. However, the inhibition of cellular contractility was more complete at the higher doses of ulacamten with IC_50_ values consistent across different species. Ulacamten demonstrated efficacy in terms of improving relaxation parameters in hypercontractile R403Q EHTs. Diastolic function also improved in ZSF1 obese rats after ulacamten treatment, with favorable outcomes for both IVRT and fibrosis.

## CLINICAL PERSPECTIVES

### COMPETENCY IN MEDICAL KNOWLEDGE

The outcome of the phase 2 AMBER-HFpEF clinical trial will determine whether ulacamten demonstrates sufficient therapeutic effects to warrant progression to a larger phase 3 study. Regardless, current data testing a variety of therapeutic candidates in patients with HFpEF point to the need for stratification of the HFpEF population based on deeper understanding of disease etiology and predictive biomarkers.^48^ Preclinical discovery of various CMIs has demonstrated there are multiple sites in myosin that can be targeted to inhibit ATPase activity in the sarcomere. Additionally, the variability in the maximal inhibition of biochemical activity across the CMIs further indicates these agents have different mechanisms of action. HFpEF is a heterogeneous disease with a hypercontractile subgroup that has overlapping features of cardiac structure and function with HCM. If CMIs are useful in treating HFpEF, there will be broader use of this class of clinical agents especially in those with high unmet medical need.

### TRANSLATIONAL OUTLOOK

The efficacy of CMIs in patients with oHCM is well established by the clinical trials of mavacamten and aficamten. However, the equivocal effect of mavacamten in a trial in patients with nHCM may indicate that a CMI may not work for all hypercontractile cardiac diseases. Ulacamten is a new class of CMI with a distinct mechanism of action from aficamten and mavacamten. Our studies of hypercontractile EHTs and a HFpEF rat model showed that ulacamten improved relaxation parameters in a dose-dependent manner. The outcome of the ongoing AMBER-HFpEF trial could further warrant the clinical development of ulacamten for a large, randomized phase 3 trial of hypercontractile HFpEF.

## HIGHLIGHTS

- CMIs proved effective in clinical trials to improve exercise capacity among oHCM patients.
- CMIs reduce excessive contractility and reverse adverse cardiac LV modeling in oHCM.
- The hypercontractile patient subgroup of HFpEF shows abnormal cardiac structure and function.
- Ulacamten is a new class of CMI with distinct properties than previous clinically proven CMIs.
- Ulacamten is being tested in a clinical trial of HFpEF with patients having EF >60%.

## DATA SHARING STATEMENT

The data underlying this article may be shared upon reasonable request to the corresponding author.

## ACKNOWLEDGEMENTS

Editorial support was provided by David Sunter, PhD, on behalf of Engage Scientific Solutions, and was funded by Cytokinetics, Inc.

## ABBREVIATIONS AND ACRONYMS

CMI: cardiac myosin inhibitor
EF: ejection fraction
EHT: engineered heart tissue
HFpEF: heart failure with preserved ejection fraction
HMM: heavy meromyosin
ITC: isothermal titration calorimetry
IVRT: isovolumic relaxation time
LV: left ventricle
RLC: regulatory light chain
S1: subfragment-1

