## Supplemental Materials for "Ulacamten: A Novel, RLC-Targeting Cardiac Myosin Inhibitor for Potential Treatment of Cardiac Hypercontractility, Including HFpEF"

**SUPPLEMENTAL APPENDIX**

**Supplemental Table 1.**	3
Concentration Dependence of Ulacamten Inhibition of Myofibrils From Different Muscle Types at pCa_75_

**Supplemental Table 2.**	4
Concentration Dependence of Ulacamten Inhibition of Actin-Activated ATPase Activity of HMM and Chymotryptic S1 Forms of Bovine Cardiac Myosin

**Supplemental Table 3.**	5
Thermodynamic Parameters for Ulacamten Binding to Recombinant Human RLC + HCF

**Supplemental Table 4.**	6
Thermodynamic Parameters for Ulacamten Binding to Bovine Cardiac Papain S1

**Supplemental Figure 1 (related to Figure 1).**	7
Ulacamten and its Inactive Enantiomer CK-4022425.

**Supplemental Figure 2 (related to Figure 1)**.	8
Specificity of Ulacamten Inhibition of Skeletal Myofibrillar ATPase Activity Versus Smooth Muscle HMM.

**Supplemental Figure 3 (related to Figure 1).**	10
Assessment of Binding of Ulacamten and CK-4022425 (Inactive Enantiomer) to Papain S1 or Multiple Fragments of Recombinant RLC.

**Supplemental Figure 4 (related to Figure 2).**	11
Functional Characterization of Contractility of Cardiomyocytes Isolated From ZSF1 Lean Rats.

**Supplemental Figure 5 (related to Figure 2)**.	12
Characterization of Baseline Contractility of EHTs Derived From Wild-Type (n=5) and R403Q iPSC Cardiomyocytes (n = 10) Demonstrating Hypercontractile Phenotype With Impaired Contraction and Relaxation Kinetics.

**Supplemental Figure 6 (related to Figure 2)**.	13
Ulacamten Dose-Response Inhibition of Wild-Type EHTs (n = 5).

**Supplemental Figure 7 (related to Figure 2)**.	14
Ulacamten Dose-Response Inhibition of R403Q EHTs (n = 10).

**SUPPLEMENTAL TABLE 1 Concentration Dependence of Ulacamten Inhibition of Myofibrils From Different Muscle Types at pCa_75_**

|  | **Cardiac** | **Slow skeletal** | **Fast skeletal** |
| --- | --- | --- | --- |
| EC_50,_ µM | 2.94 (2.74-3.16) | 2.51 (1.65-4.64) | 11.9 (9.3-17.8) |
| IC_15,_ µM | 1.43 (1.35-1.50) | 1.42 (1.11-1.78) | 3.31 (2.94-3.70) |
| Hill slope | −1.19 (−1.29 to −1.09) | −0.94 (−1.4 to −0.52) | −1.13 (−1.36 to −0.91) |
| Top | 1.01 (1.00-1.02) | 1.05 (1.01-1.16) | 0.99 (0.98-1.01) |
| Bottom | 0.47 (0.46-0.49) | 0.50 (0.33-0.57) | 0.24 (0.085-0.33) |

Values are the output from nonlinear least squares fitting of ATPase reactions in Figure 1A and Supplemental Figure 1C.

The 95% confidence interval is shown in parentheses.

EC_50_ = the ulacamten concentration for 50% of the maximal inhibition; IC_15_ = the ulacamten concentration at which 15% of inhibition; pCa_75_ = 75% of maximum calcium ion activation; Top and Bottom are the top and bottom plateaus in the unit of Y.

**SUPPLEMENTAL TABLE 2 Concentration Dependence of Ulacamten Inhibition of Actin-Activated ATPase Activity of HMM and Chymotryptic S1 Forms of Bovine Cardiac Myosin**

|  | **HMM** | **S1** |
| --- | --- | --- |
| EC_50,_ µM | 2.54 (2.32-2.79) | NA |
| IC_15,_ µM | 1.03 (0.97 to 1.08) | >39 |
| Hill slope | −0.87 (−0.95 to −0.79) | NA |
| Top | 1.01 (1.00-1.03) | NA |
| Bottom | 0.49 (0.47-0.51) | NA |

Values are the output from nonlinear least squares fitting of ATPase reactions in Figure 1C.

The 95% confidence interval is shown in parentheses.

EC_50_ = the ulacamten concentration for 50% of the maximal inhibition; HMM = heavy meromyosin; IC_15_ = the ulacamten concentration at which 15% of inhibition; NA = not available; S1 = subfragment-1; Top and Bottom are the top and bottom plateaus in the unit of Y.

**SUPPLEMENTAL TABLE 3 Thermodynamic Parameters for Ulacamten Binding to Recombinant Human RLC + HCF**

|  | **RLC + 12-hep HCF**  **(n = 10)** | **RLC + 2-hep HCF**  **(n = 12)** |
| --- | --- | --- |
| Stoichiometry, N^a^ | 0.78 ± 0.10 | 0.94 ± 0.084 |
| Affinity, KD, µM | 7.3 ± 1.3 | 6.6 ± 1.5 |
| Enthalpy, ΔH, kcal/mol | −6.1 ± 1.2 | −5.3 ± 0.65 |
| Entropy, ΔS, e.u. | 3.0 ± 4.1 | 6.0 ± 2.6 |

Data shown are mean ± standard deviation. Representative binding isotherms are shown in Figure 1E.

^a^Stoichiometry is expressed relative to the concentration of RLC-containing monomers

HCF = heavy chain fragments; hep = heptads; RLC = regulatory light chain

**SUPPLEMENTAL TABLE 4 Thermodynamic Parameters for Ulacamten Binding to Bovine Cardiac Papain S1**

|  | **Bovine Cardiac Papain S1**  **(n = 6)** |
| --- | --- |
| Stoichiometry, N | 0.54 ± 0.04 |
| Affinity, KD, µM | 3.9 ± 1.0 |
| Enthalpy, ΔH, kcal/mol | -5.7 ± 0.4 |
| Entropy, ΔS, e.u. | 5.8 ± 1.7 |

Data shown are mean ± standard deviation

Representative binding isotherms are shown in Supplemental Figure 3A.

**SUPPLEMENTAL FIGURE 1** **Ulacamten and its Inactive Diastereomer CK-4022425**. The chemical structures of ulacamten and its negative control CK-4022425 are shown. CK-4022425 has no inhibitory effect on bovine cardiac myofibrillar ATPase activity. ATPase activities were measured at the Ca^2+^ concentration of 75% activation. Data represent mean ± standard deviation.

**Screening hit 🡪**Ulacamten

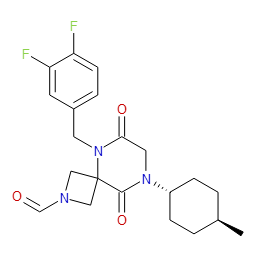

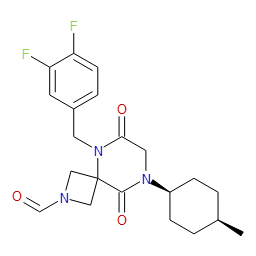

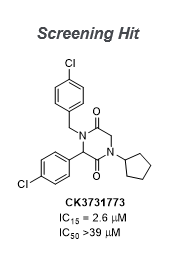

CK-4022425

(neg. Control)

🡪 🡪 🡪

Screening Hit

Ulacamten

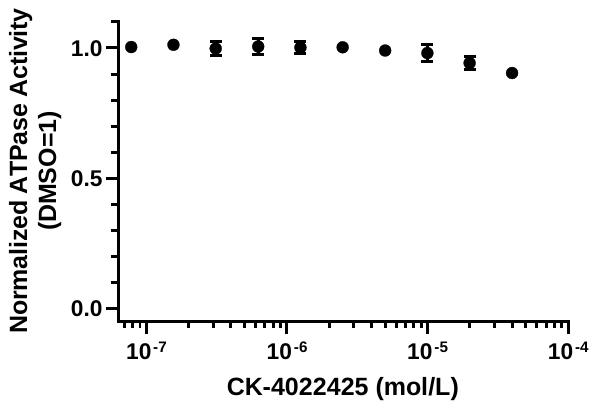

Myofibril ATPase dose-response for CK-4022425

Ca^2+^ = calcium ion-positive

**SUPPLEMENTAL FIGURE 2** **Specificity of Ulacamten Inhibition of Skeletal Myofibrillar ATPase Activity Versus Smooth Muscle HMM.** (A) Inhibition of myosin ATPase activity in slow (square) and fast (inverse triangles) skeletal myofibrils by ulacamten. ATPase activities were measured at the Ca^2+^ concentration of 75% activation. Data represent mean ± SD with 4-parameter fitting. The biochemical EC_50_ which represents the ulacamten concentration for 50% of the maximal inhibition was 2.5 μM (n = 8; 95% CI 1.65-4.64) and 11.9 μM (n = 20; 95% CI 9.3-17.8). (B) Testing inhibition of myosin ATPase activity in smooth muscle HMM by ulacamten. Dose-response curves of actin-activated myosin ATPase activity are shown. Data represent mean ± SD with 4-parameter fittings. (C) Testing myosin inhibition in single-headed papain S1 by ulacamten. Dose-response curves of actin-activated myosin ATPase activity are shown. Data represent mean ± SD with 4-parameter fittings. (D) SDS PAGE gel of myofibrils before and after RLC depletion. Lane 3 and Lane 4 represent myofibrils after and before RLC depletion, respectively. Lane 1 represents full-length myosin, which had myosin heavy chain, essential light chain (ELC), and RLC as indicated by arrows. Full length myosin helped to find the band for RLC in myofibrils which had various sarcomeric proteins. Lane 2 represents protein standards.

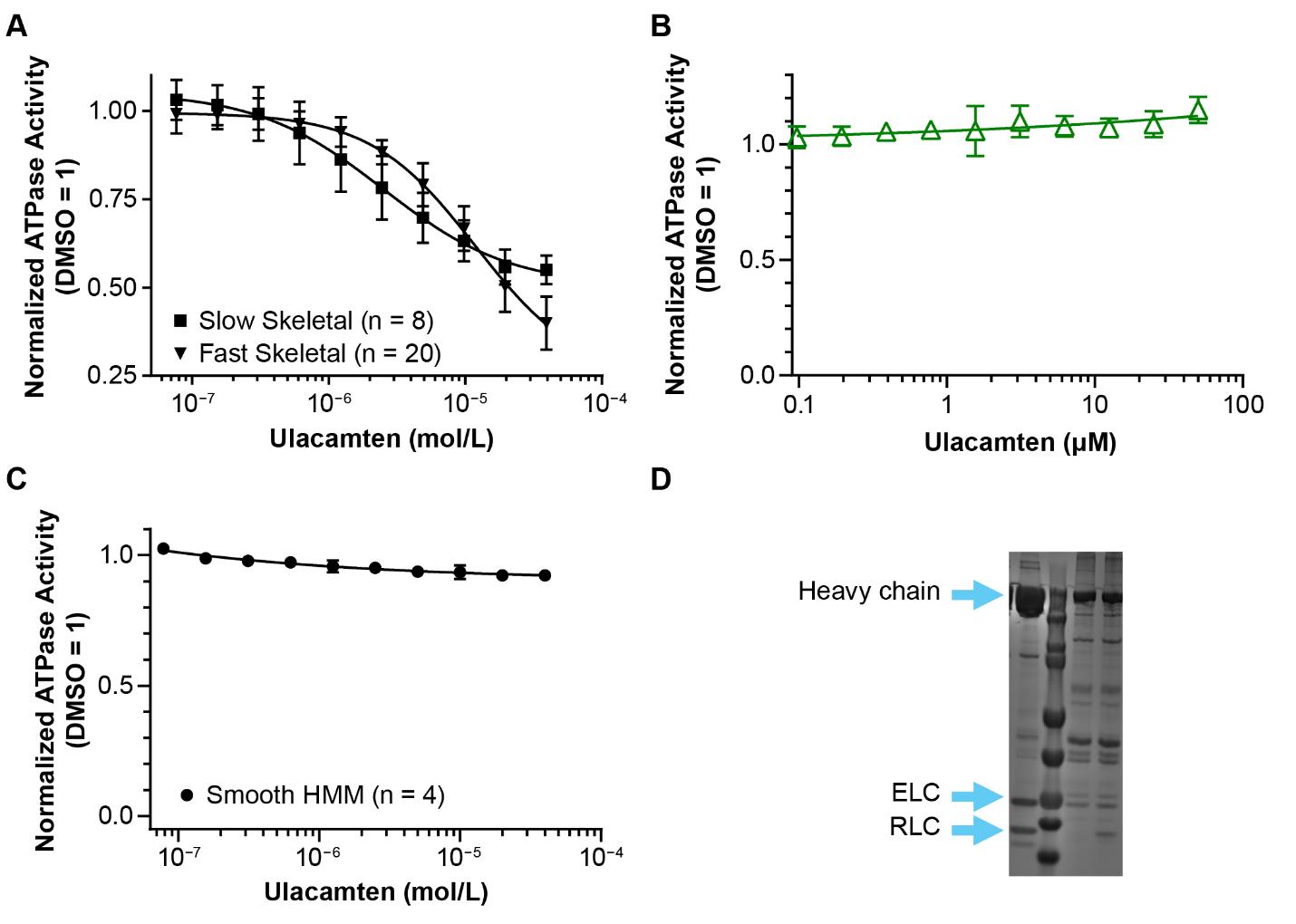

Ca^2+^ = calcium ion-positive; CI = confidence interval; EC_50_ = half-maximal concentration of biochemical inhibition; HMM = heavy meromyosin; RLC = regulatory light chain; S1 = subfragment-1; SD standard deviation

**SUPPLEMENTAL FIGURE 3** **Assessment of Binding of Ulacamten and CK-4022425 (Inactive Enantiomer)** **to Papain S1 or Multiple Fragments of Recombinant RLC.** (A) Ulacamten but not CK-4022425 binds to papain S1. (B) CK-4022425 does not bind to recombinant myosin HCF.

**A**

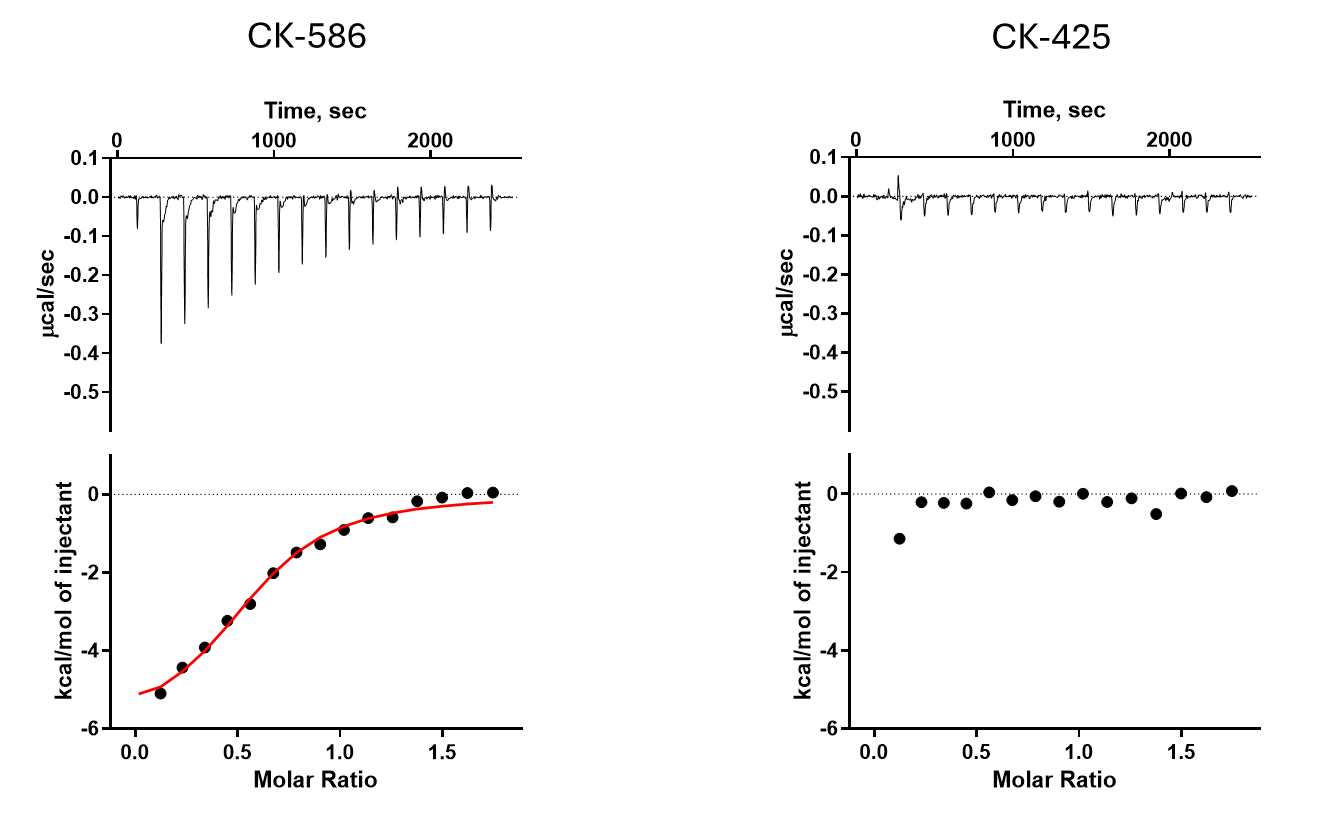

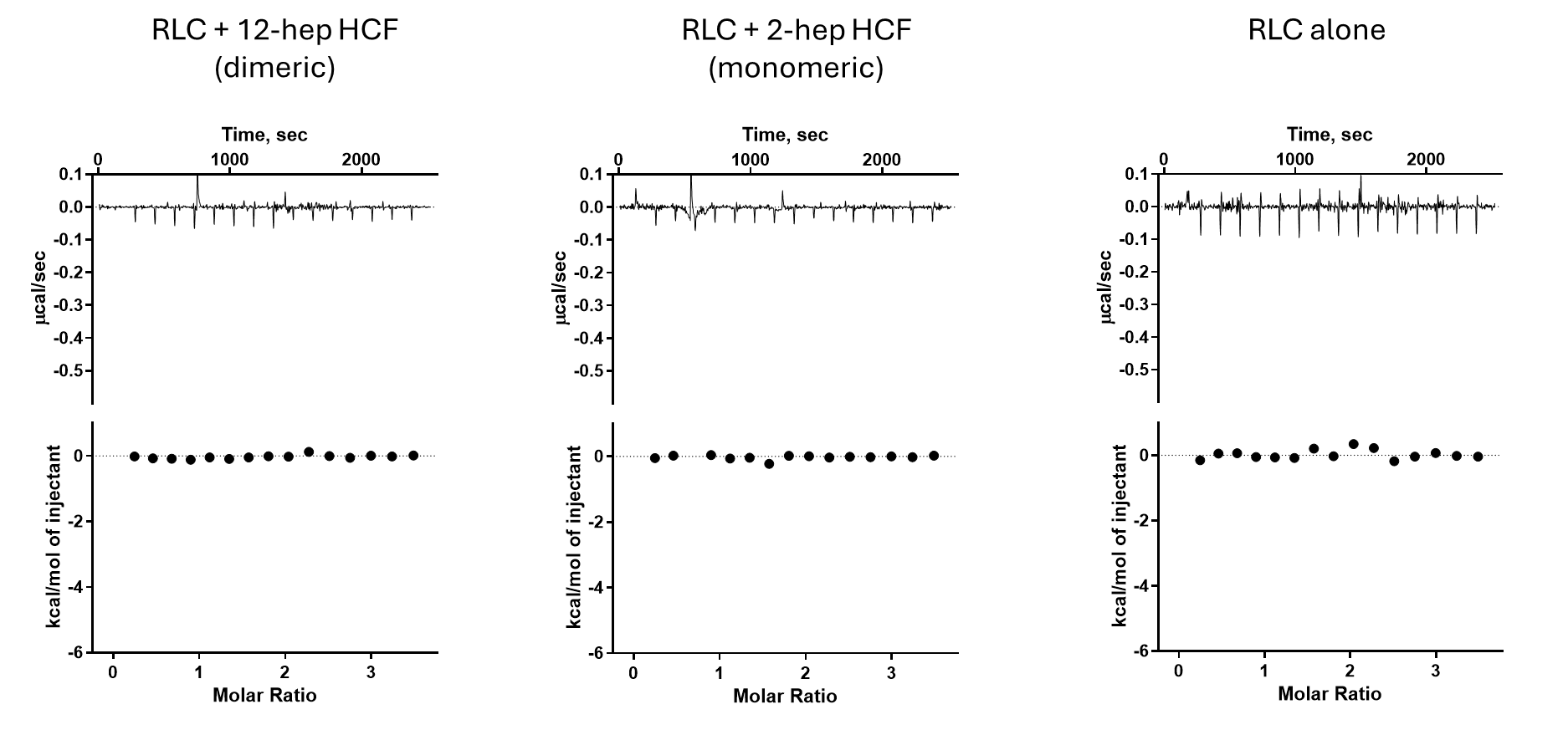

ITC data for binding to papain S1

ITC data for binding of CK-4022425 (neg control)

**B**

HCF, heavy chain fragment; ITC, isothermal titration calorimetry; RCL, regulatory light chain; S1, subfragment-1.

**SUPPLEMENTAL FIGURE 4** **Functional Characterization of Contractility of Cardiomyocytes Isolated From ZSF1 Lean Rats.** (A) Analysis of Fura-2 calcium transients in isolated ZSF1 lean rat myocytes. Values for each parameter are presented as response from baseline following treatment with 10 µM ulacamten or vehicle control (N = 3 isolations, >35 cells). Statistical significance was evaluated with unpaired Welch’s student’s t-test assuming unequal variance. (B) Sarcomere length fractional shortening (normalized to baseline) with ulacamten dose-response inhibition in ZSF1 lean rat myocytes (N = 3 isolations, >30 cells). IC_50_ and Hill slope were calculated using 4-parameter logistic dose response fit.

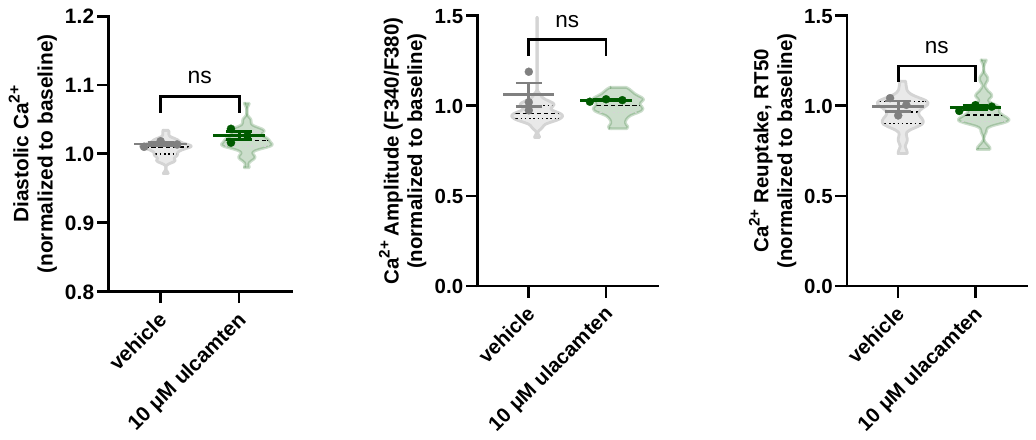

**A**

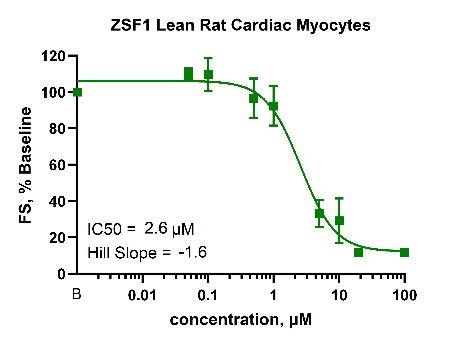

**B**

FS = fractional shortening; IC_50_ = half-maximal inhibitory concentration; ns = not significant

**SUPPLEMENTAL FIGURE 5** **Characterization of Baseline Contractility of EHTs Derived From Wild-Type (n = 5) and R403Q iPSC Cardiomyocytes (n = 10) Demonstrating Hypercontractile Phenotype With Impaired Contraction and Relaxation Kinetics.** (A) Average force (µN), (B) time to peak force (ms), (C) return time to 90% baseline (RT_90_, ms) and (D) return time to 50% baseline (RT_50_, ms) are shown for both W-T (blue) and R403Q (green). All data is expressed as mean ± standard deviation. Significance was determined with Welch’s students t-test assuming unequal variance. ***P* < 0.01, ****P* < 0.001.

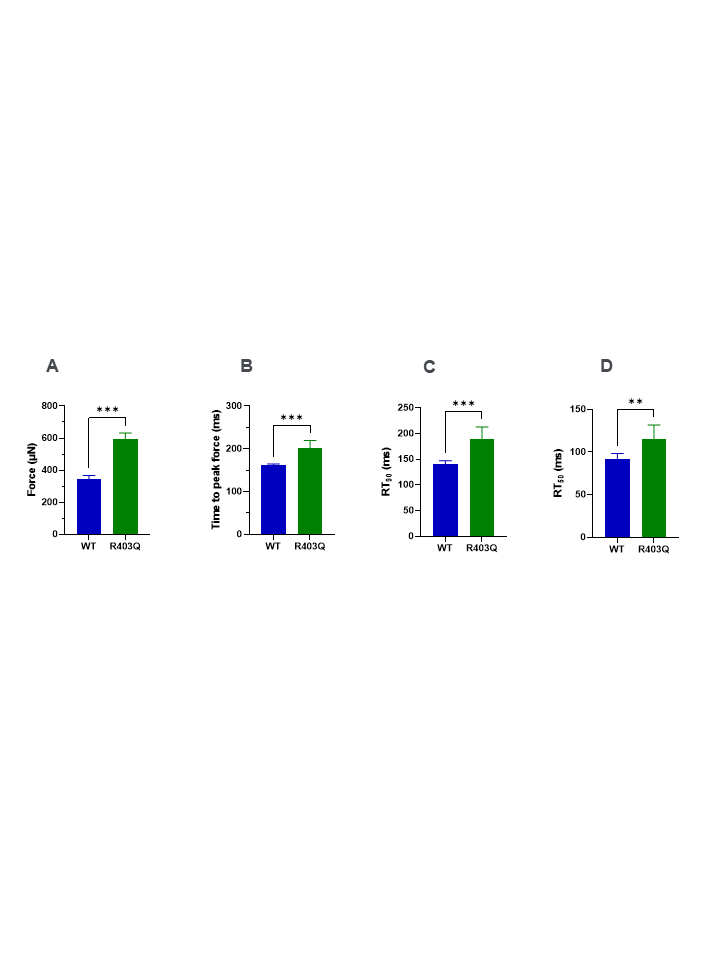

RT_50_ = return time to 50%; RT_90_ = return time to 90%; WT = wild-type.

**SUPPLEMENTAL FIGURE 6** **Ulacamten Dose-Response Inhibition of Wild-Type EHTs (n = 5).** (A) Force (µN) (B) Time to peak force (ms), (C) return time to 50% baseline, and (D) return time to 90% baseline. Solid and dotted lines represent the values for untreated and their standard deviations, respectively.

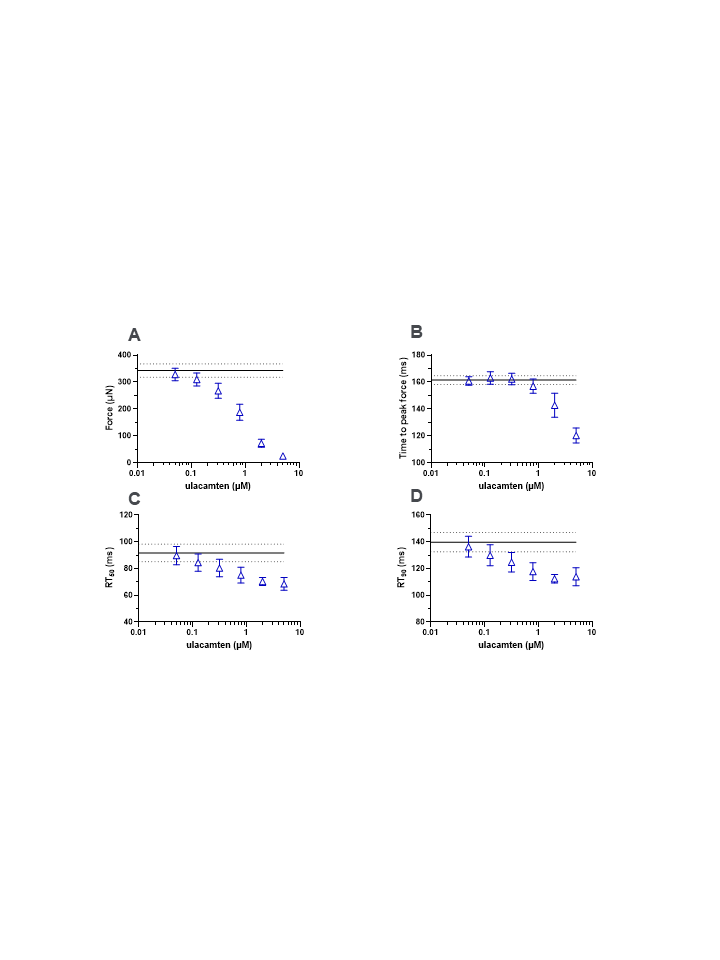

EHT = engineered heart tissue; RT_50_ = return time to 50%; RT_90_ = return time to 90%.

**SUPPLEMENTAL FIGURE 7** **Ulacamten Dose-Response Inhibition of R403Q EHTs (n = 10).** (A) Force (µN) (B) Time to peak force (ms), and (C) return time to 90% baseline. Solid and dotted lines represent the values for WT and their standard deviations, respectively.

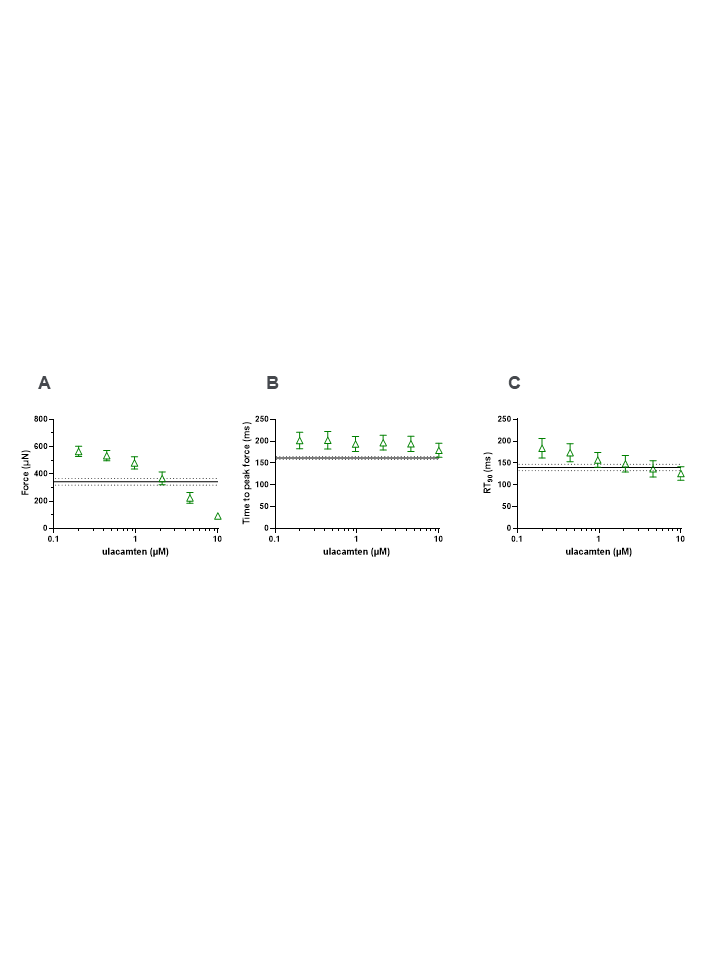

EHT = engineered heart tissue; RT_90_ = return time to 90%.
